# Empirical Estimation of Ambient Contamination in Combinatorial Single-Cell Methods Using Multi-Reference Mapping

**DOI:** 10.64898/2026.09.11.750809

**Authors:** Fabio Gomez-Cano, Luguang Jiang, Joshua D. Welch, Alexandre P. Marand

## Abstract

Droplet-based microfluidics and combinatorial indexing (scifi-ATAC and scifi-RNA) have made single-cell experiments massively scalable. However, higher-order multiplexing complicates data quality, introduces noise, and affects the potential for biological discovery. Here, we show that ambient chromatin accumulates through the experimental workflow and distorts chromatin profiles, most drastically in low-depth nuclei and in minority populations. Standard cell calling relies heavily on read count thresholds, while existing decontamination methods generally operate on aggregated count matrices rather than the underlying reads. We introduce scifi-demux, for preprocessing scifi-ATAC libraries, and AmbientMapper, a generative model that maps reads competitively against multiple references, learns the ambient profile from empty and low-complexity barcodes, and separates nuclei from background and singlets from doublets by Bayesian Information Criterion. Using interspecies ground truth experiments, published multi-genotype libraries, and simulated read-level synthetic barcodes in which every contaminating read is traceable, we show that calls are robust to parameter variation and stable across designs, achieving a wrong-genome rate of 0.19% on a 26-genome reference panel. Finally, we evaluated the impact of removing contaminants, showcasing how AmbientMapper rescues low-depth nuclei discarded by standard pipelines and restores biological structure obscured by contamination.

## INTRODUCTION

The widespread adoption of droplet-based microfluidics^1,2^ and the development of combinatorial indexing^3–5^ have exponentially increased the scale of single-cell experiments^6^. However, as cell profiling scales, so does the introduction of systematic noise, widely recognized as ambient contamination^7,8^. In droplet-based assays, this phenomenon can be driven by the encapsulation of cell-free molecules that distort single-cell molecular profiles, inflating apparent multiplet rates, and confounding downstream tasks such as clustering and differential analysis^8–10^. While combinatorial indexing strategies are theoretically less susceptible to such noise due to multi-step barcoding, this assumption holds only if nuclear integrity is perfectly preserved. However, complex split-pool and plate-indexing workflow introduces unique vulnerabilities, including ambient molecules accumulation from sample processing and technical artifacts such as index hopping and PCR chimeras. Consequently, the noise landscape of combinatorial indexing remains poorly characterized. Accurately resolving these artifacts requires a framework capable of discerning true cellular signals from background noise and precisely assigning the genomic origin of each sequence read.

A rich ecosystem of bioinformatics tools has been developed to address demultiplexing, genotyping, and contamination estimation. These methods generally fall into three broad categories. Genetic demultiplexers, such as Demuxlet^11^, Vireo^12^, scSplit^13^, Souporcell^14^, and Demuxalot^15^ assign barcodes to specific donors based on genotypic variation. While these tools incorporate explicit doublet models, they implicitly treat contaminated cells as genetic mixtures rather than resolving the background noise itself. Tag-based methods, including Cell Hashing^16^ and DemuxEM^17^, utilize experimental priors to separate signal from background via mixture modeling. Finally, ambient estimators like EmptyDrops^18^, SoupX^7^, DecontX^19^, and CellBender^8^ focus on modeling the background profile, typically derived from cell-free droplets, to probabilistically denoise count matrices. Recent work has moved further in this direction, with Ambimux classifying droplets and estimating per-modality ambient fractions from donor genotypes at variant sites^20^, and CellSweep resolving observed counts into cell-type, ambient, and global bulk components under a multinomial mixture^21^. Despite this methodological breadth, a critical gap remains. No unified framework exists that (i) operates directly at the read level across multiple reference genomes, (ii) models ambient contamination as a competing origin for each individual read, weighed against every candidate reference genome, (iii) performs cell-level model selection without variant calls, optionally guided by the combinatorial design of the library, and (iv) systematically evaluates the impact of these artifacts on combinatorial indexing assays. These gaps are particularly limiting for experiments involving large cohorts of pooled genotypes.

Here we introduce AmbientMapper, a generative probabilistic framework designed to resolve ambient contamination and cellular identity in combinatorial indexing assays. We hypothesized that the primary source of contamination in these protocols is not random sequencing noise, but cell-free chromatin released from compromised nuclei during the initial indexing steps, which then co-segregate with intact nuclei. To evaluate this possibility, we designed a controlled interspecies (barnyard) scifi-ATAC-seq experiment using *Zea mays* (maize) and *Arabidopsis thaliana* (Arabidopsis), where species-specific nuclei were indexed in separate wells. Using this barnyard design, we demonstrate how competitive mapping to multiple reference genomes enables the precise detection of ambient contamination, quantification of cross-plate contamination, and discrimination of true biological doublets from low-level background. We then test the framework where its assumptions are hardest to satisfy, using read-level simulations in which every contaminating read is traceable, on a single-genotype library mapped against 26 distinct genomes, and finally, against an orthogonal variant-based demultiplexer on two multi-genotype libraries. While demonstrated here on scifi-ATAC-seq, the principles of AmbientMapper extend naturally to scifi-RNA-seq and standard droplet-based workflows where multiple-genome references are part of the experimental design, providing a robust solution for denoising population-scale high-throughput single-cell datasets. Alongside AmbientMapper, we describe scifi-demux, a preprocessing tool that performs plate and well demultiplexing, barcode correction and adapter handling that scifi-ATAC-seq libraries require before any analysis is possible.

## RESULTS

### Ambient chromatin contamination in combinatorial indexing assays is pervasive, asymmetric, and confounding

We hypothesized that ambient contamination in combinatorial indexing (scifi-ATAC-seq) protocols originates from nuclear compromise during sample handling and indexing, which releases a pool of cell-free chromatin that acquires a well index after the wells are pooled and then co-segregates with intact nuclei into droplets (**Fig. 1A**). To test this hypothesis under controlled conditions, we designed an interspecies scifi-ATAC-seq experiment using maize and Arabidopsis nuclei, indexed in separate wells on a 96-well plate (**Table S1, S2**). This design enables unambiguous discrimination between true cellular signal and cross-species ambient noise based on physical plate coordinates rather than post hoc inference. Following demultiplexing and competitive alignment to individual reference genomes, we genotyped reads to assess barcode purity. While most barcodes mapped predominantly to their expected genome, we observed pervasive cross-species signals (**Fig. 1B**). Quantifying off-target read fractions revealed that contamination was strongly asymmetric. The mean off-target fraction was 0.73 in Arabidopsis-indexed barcodes compared to 0.04 in maize-indexed barcodes (**Fig. 1C**, **1D**). Additionally, whereas 84% of maize barcodes contained less than 1% off-target reads, only 8% of Arabidopsis barcodes reached comparable purity. Conversely, 67% of Arabidopsis barcodes carried at least 64% maize-derived reads, against 2.7% of maize barcodes at the same threshold (**Fig. 1D**). This asymmetry was robust to mapping strategy: re-analysis using a concatenated reference genome yielded the same systematic bias, ruling out alignment artifacts as the primary driver (**Fig. S1**). This bias also cannot be explained by shallow barcode coverage. Among barcodes passing standard nucleus quality control, the off-target fraction of Arabidopsis-indexed barcodes increased to 0.93 compared to 0.04 in maize, implying that nucleus-like barcodes are the most contaminated. These results indicate that ambient contamination is structured and driven by the dominant chromatin mass in the pool, which in this case was the larger maize genome (maize genome ∼ 2.2Gb vs. Arabidopsis genome ∼ 0.135Gb)^22,23^. Asymmetric contamination could also be exacerbated by the different nuclei isolation optima of each species. For example, Arabidopsis nuclei require lower concentrations of detergents to maintain nuclear integrity (See *Methods*). Consistent with this model, analysis of dominance ratios (top-1 versus top-2 genome assignments) showed that contamination effects are most pronounced in low-depth barcodes (<1,000 reads), creating a “ratio trap” in which true low-coverage nuclei become statistically indistinguishable from ambient signal (**Fig. 1E**).

**Fig. 1.**
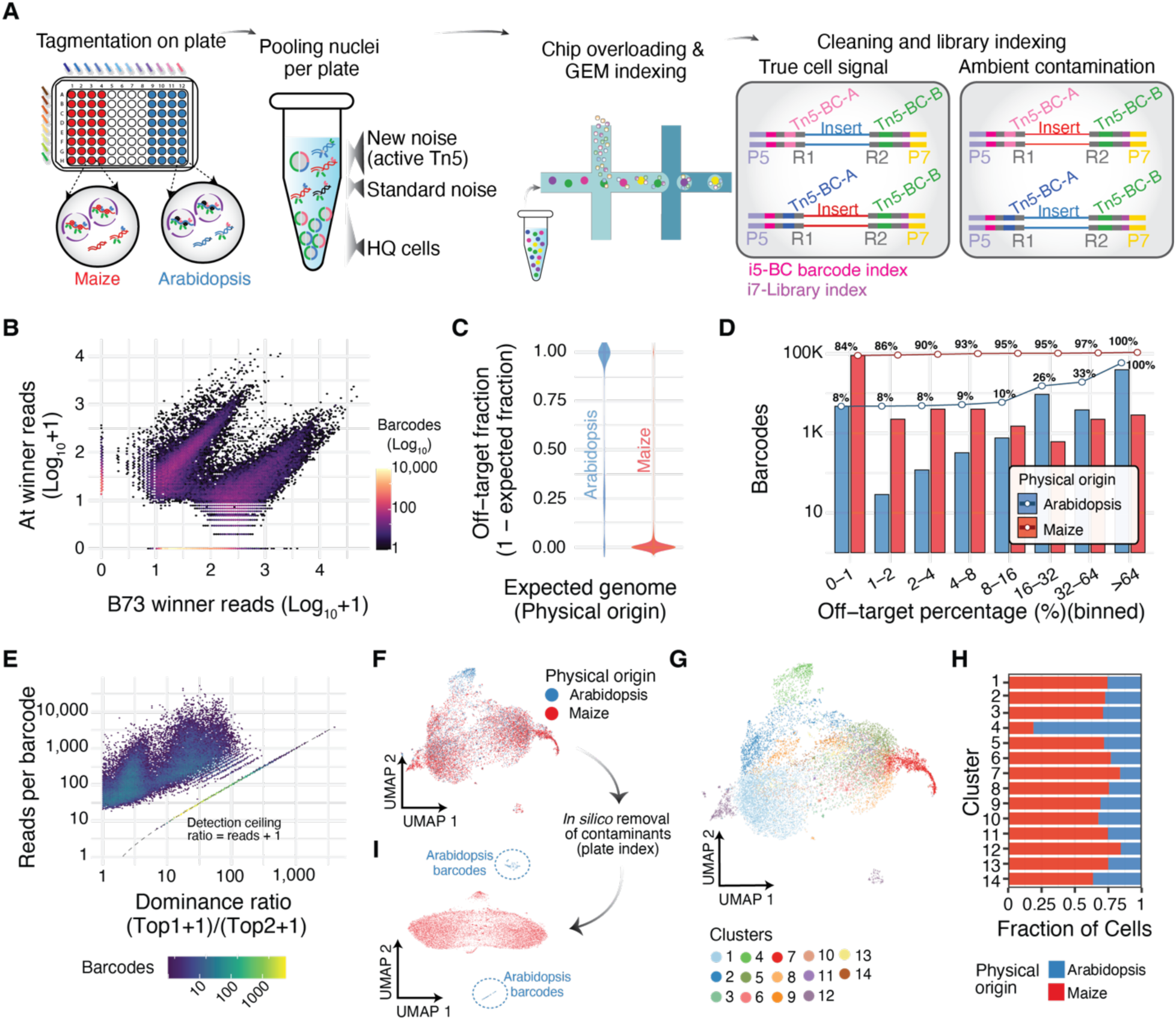
Pervasive and asymmetric ambient contamination in combinatorial scifi-ATAC-seq assays. **A,** Schematic of the scifi-ATAC-seq design and proposed origin of ambient contamination. Maize and *Arabidopsis* nuclei were loaded into separate wells of a 96-well plate, such that each Tn5 index identifies the species loaded into that well. Chromatin released from compromised nuclei can be tagmented after pooling by residual active Tn5, acquire an index inconsistent with its species of origin, and subsequently be co-encapsulated with intact nuclei. **B,** Barnyard density plot showing unique reads mapping to *Zea mays* (B73) versus *Arabidopsis thaliana* for each barcode. Reads were assigned to the genome with the highest alignment score (“winner reads”). **C,** Distribution of the off-target read fraction for barcodes indexed in *Arabidopsis* or maize wells. Contamination was strongly asymmetric, with *Arabidopsis*-indexed barcodes containing a larger maize-derived fraction. **D,** Distribution of barcodes across contamination-rate bins, stratified by plate origin. Lines show the cumulative fraction of barcodes at each threshold. **E,** Density plot of the pseudocount dominance ratio, (k₁ + 1)/(k₂ + 1), where k₁ and k₂ denote reads assigned to the highest- and second-ranked genomes, respectively. Low-depth barcodes frequently showed weak dominance ratios that overlapped with background. **F,** UMAP of all barcodes based on chromatin accessibility after independent mapping to each reference genome and joint embedding, colored by expected genome from the plate index. **G,** Same UMAP colored by Leiden cluster. **H,** Cluster purity, showing the proportion of *Arabidopsis*- and maize-indexed barcodes within each Leiden cluster. **I,** UMAP after in silico removal of cross-genome reads, using the plate index as ground truth so that each barcode retains only reads from the genome loaded into its well.

To evaluate the impact of ambient contamination on downstream data quality, we summarized quality metrics for each plate population with at least 200 reads (**Fig. S2**). Arabidopsis barcodes exhibited a median fraction of reads in peak (FRiP) of 0.61 compared to 0.68 in maize. The median fraction of reads within 1-kb of transcription start sites (TSSs) was also similar (0.47 in Arabidopsis and 0.45 in maize; **Fig. S2**). Given that all these values fall within ranges typically associated with high-quality scATAC-seq data, we interpret this as indicating that standard QC filters alone cannot detect ambient contamination. Contamination was also evident in the low-dimensional embedding (**Fig. 1F**) and within shared Leiden communities (**Fig. 1G**, **1H**). Importantly, this co-embedding (**Fig. 1F**) was built by mapping each barcode to each genome independently and merging the count matrices *in silico*. Consequently, two barcodes are placed together only when their reads fall in the same regions of the same genomes. Therefore, co-projection of maize and Arabidopsis barcodes is indicative of cross-species mapping rather than cellular similarity. Consistently, removing cross-genome reads *in silico* results in cleanly separated species and reduced mixing from near-random levels to zero (**Fig. 1I**). Co-projection of maize and Arabidopsis barcodes could also result from loss of high dimensional relationships artificially because of UMAP parameter choices. To exclude this possibility, we evaluated embeddings across a grid of heuristic UMAP parameters. For each configuration, we quantified neighborhood preservation relative to the Principal Component Analysis (PCA) space, genome mixing within local neighborhoods, and correlations between embedding coordinates and QC covariates. Across all tested parameter combinations, genome mixing remained stable and correlations with QC covariates remained low (**Fig. S3**). Arabidopsis and maize barcodes stayed mixed regardless of how well the embedding preserved the PCA neighborhoods, indicating that the observed mixing is a property of the reads in each barcode and not the embedding parameters. Collectively, these results demonstrate that ambient chromatin contamination is a systematic artifact that (i) overwhelmingly impacts the minority population, (ii) biases standard quality control, and (iii) introduces structured technical variation that survives conventional quality control and propagates into dimensionality reduction and clustering (**Fig. 1F**-**1H**), where it can be misinterpreted as a biological signal.

### A probabilistic framework for resolving cellular identity from ambient noise in genomic mixing experiments

Combinatorial indexing assays increasingly leverage genomic multiplexing, pooling distinct species or genotypes within a single experiment. Beyond increasing throughput, pooling also provides an empirical estimate of the multiplet rate and reduces batch effects. However, as shown above, ambient chromatin compromises such datasets most severely at low coverage and in minority populations, where true signal becomes statistically indistinguishable from background noise. To address this challenge, we developed AmbientMapper, a generative probabilistic framework that explicitly models sequenced reads as a mixture of true cellular signal and ambient background (**Fig. 2A**). In contrast to standard pipelines that rely on fixed read-count thresholds or post hoc filtering, AmbientMapper operates at read-level resolution and learns experiment-specific noise structure directly from the data. The workflow begins with a competitive alignment against every candidate reference genome assembly. For each read *r*, the alignment evidence across genomes is summarized using three metrics: alignment score, mapping quality, and edit distance, which are then calibrated against empirical cumulative distribution (ECDFs). Then, each metric is fused into a per-genome posterior likelihood (*L_r_*_,#_), for read *r* originated from genome *g* (**Fig. S4**). Barcodes with low complexity are treated not as sequencing failures but as an empirical estimate of the ambient profile, η. This profile is refined iteratively by retaining candidate barcodes whose read composition is closest to the current estimate, as measured by Jensen Shannon divergence.

**Fig. 2.**
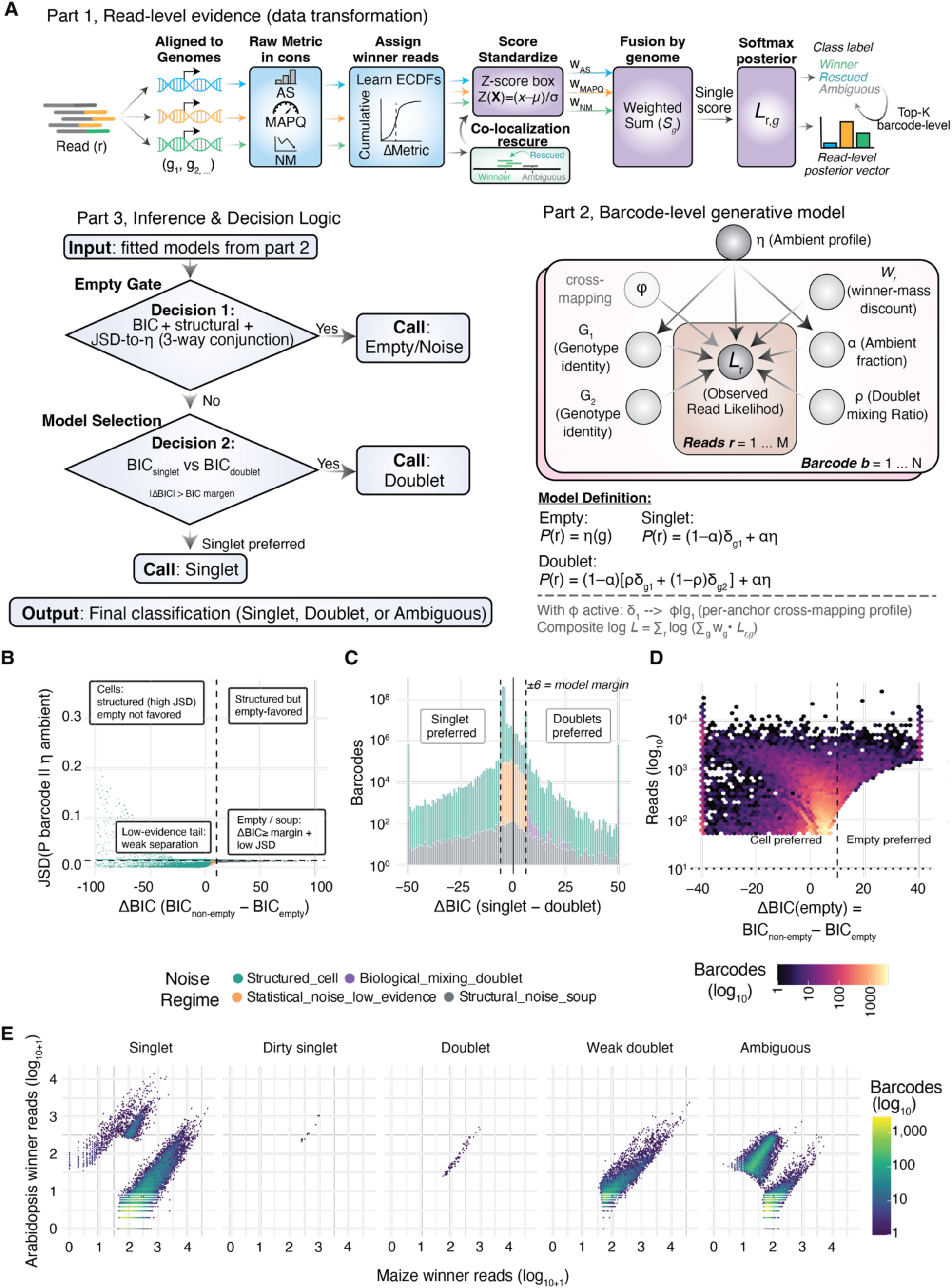
AmbientMapper: a generative probabilistic framework for genotyping and ambient removal. **A,** Schematic of the AmbientMapper workflow. *Read-level evidence:* reads are competitively aligned to all candidate reference genomes, alignment metrics are calibrated using depth-stratified empirical cumulative distribution functions (ECDFs), and combined into a per-genome likelihood, **L**(r,g), and read-reliability weight, wᵣ. *Barcode-level model:* weighted reads are evaluated under empty, singlet, and doublet hypotheses using an ambient profile, η, learned from low-depth barcodes, followed by BIC-based model comparison. *Inference:* ambient-like barcodes are rejected using the empty-model BIC advantage together with structural diffuseness and, optionally, divergence from η (Decision 1). Remaining barcodes are classified as singlets, doublets, or intermediate call classes using the singlet–doublet ΔBIC and purity/strength criteria (Decision 2). Read-level calibration and parameter values are detailed in Fig. S4 and Methods. **B,** Empty-barcode classification. Jensen–Shannon divergence (JSD) from η plotted against the ΔBIC favoring the empty model, colored by inferred noise regime. The vertical dashed line indicates the empty BIC-margin threshold. Barcodes are classified as empty only when the BIC and structural-diffuseness criteria are jointly satisfied. The horizontal dashed line denotes τ, used only to visualize the ambient-like regime and not for classification. **C,** Singlet- versus-doublet model selection for barcodes passing the empty gate. Distribution of ΔBIC, with dashed lines at ± bic_margin delimiting the ambiguous region. **D,** Read depth versus ΔBIC favoring the empty model. Discrimination increases with coverage, producing a funnel-shaped distribution in which low-depth barcodes concentrate near zero and higher-depth barcodes separate into cell-and empty-preferred populations. **E,** Barnyard plots stratified by AmbientMapper call, showing the principal *singlet*, *doublet*, *dirty singlet*, and *weak doublet* classes and their distinct read-composition profiles. Full class definitions and sub-gate thresholds are provided in the main text and Methods.

Barcode identity is then decided in two stages, both by Bayesian model selection under a complexity penalty. The first stage asks whether a barcode is better explained by the ambient profile alone than by any model containing a nucleus. We present this comparison as ΔBIC_empty = BIC(best non-empty model) − BIC(empty model), so that positive values favor the empty model, because Bayesian information criterion (BIC) is a penalized loss for which the lower value indicates the better fit. A barcode is called empty only when two conditions are satisfied, ΔBIC_empty above a margin of 6, and genuinely diffuse read mass. Here, we defined diffuse read mass as cases in which no genome holds more than 60% of the reads and the top genome exceeds the runner-up by no more than two-fold (Fig. 2B). The second stage classifies the barcodes that pass this gate by evaluating ΔBIC_call = BIC(singlet) − BIC(doublet), for which positive values favor the doublet model, calling a barcode singlet or doublet only when the two models differ by more than 6 BIC units and ambiguous otherwise (Fig. 2C, Fig. S4). We used 6 BIC units for both margins following the conventional interpretation of BIC differences, in which two to six units constitute positive evidence and more than six constitute strong evidence of better fit24. The two comparisons are distinct signed measures, and both inherit the depth dependence of the underlying evidence. Low-depth barcodes cluster near zero, where the empty and nucleus-bearing models are statistically indistinguishable, whereas high-depth barcodes separate into clearly cell-preferred and clearly empty-preferred populations (Fig. 2D). Thus, instead of forcing low-confidence barcodes into discrete binary classes, AmbientMapper reports a granular classification that reflects model confidence.

To evaluate this decision system, we ran AmbientMapper on the interspecies dataset and stratified the cross-species barnyard plot by the resulting calls (**Fig. 2E**). Because the plot places every barcode by its reads on the two genomes, the same evidence the model scores, each call should occupy its own region of the plot, and this is what we observed. Specifically, a “singlet” is a barcode the model is confident contains a single nucleus, in practice one dominated by a single genome, with at least 60% of the read mass on the top genome and a 1.5-fold margin over the runner-up. Most barcodes in this class therefore still carry a small fraction of reads from the other genome(s). These barcodes are flagged at the read level and removed during the design-aware decontamination step described below. Purity is therefore restored downstream, rather than by tightening the call at this stage. “Doublet” barcodes form a clear diagonal population, representing nuclear collisions. Critically, the model also identifies intermediate states, including “dirty singlet”, which maps primarily to one genome but carries enough off-target signal to fail the purity sub-gates, and “weak doublet”, which the model fits better as two components but whose secondary genome carries less than 20% of the read mass, which is the floor for a confident doublet call. This probabilistic stratification allows users to distinguish between high-quality nuclei and those affected by heavy contamination or technical artifacts, enabling principled decisions about data retention. Notably, the empty gate is conservative by construction, because it requires diffuse read mass as well as decisive evidence. Shallow barcodes are set aside by a minimum depth of five reads, and ambient signal is removed at the read level by the decontamination step described below rather than by discarding barcodes at the calling stage.

### Design-aware decontamination surgically removes ambient noise and rescues misclassified cells

Following probabilistic genotyping, AmbientMapper performs targeted decontamination to generate a clean read set for downstream analysis. By leveraging the combinatorial indexing experimental design (plate layout, e.g., **Fig 1A**) as a prior, the algorithm identifies and removes reads originating from off-target genomes while preserving signal from the expected genotype. To evaluate this approach, we compared barcode-level read distributions before and after cleaning (**Fig. 3A**). This analysis showed that AmbientMapper removes the off-diagonal contaminant signal while retaining affected barcodes for downstream quality control. This result was corroborated by the genome-wide contamination profile, which shifted from a broad, high-background distribution to a near-zero off-target fraction (**Fig. 3B**).

**Fig. 3.**
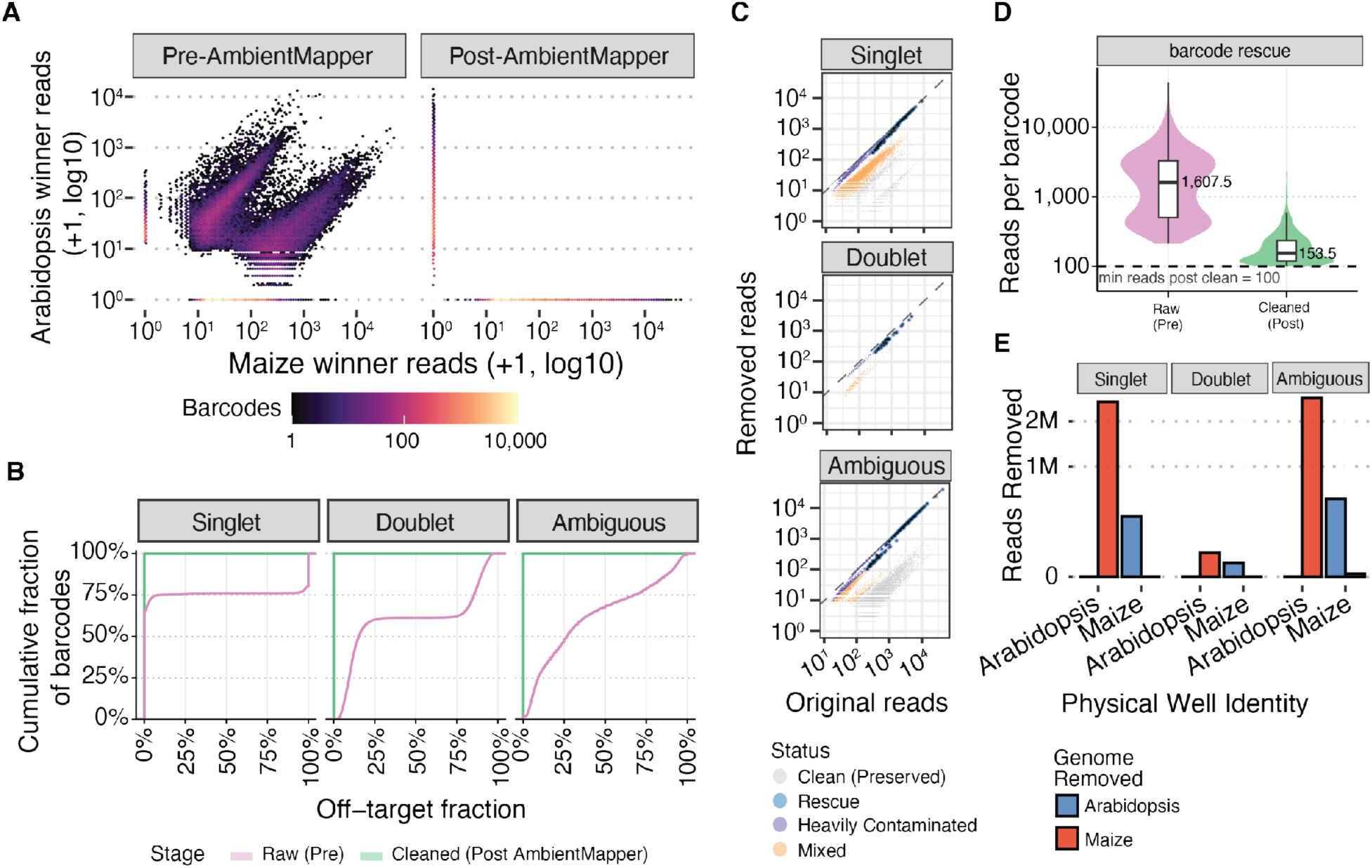
Design-aware decontamination restores library purity and resolves technical artifacts. **A,** Barnyard plots of per-barcode read counts before (left) and after (right) AmbientMapper decontamination. **B,** Empirical cumulative distributions of the per-barcode off-target read fraction (1 − expected-genome fraction), stratified by AmbientMapper call class. Before cleaning, all classes showed broad off-target distributions, including 20.1% of singlet barcodes with >99% off-target reads. After cleaning, distributions collapsed to zero, as expected when only reads assigned to the design-expected genome are retained. **C,** Reads removed per barcode versus pre-clean read depth for barcodes with >10 reads, shown on log10 scales and faceted by AmbientMapper call class. Points are colored by cleaning outcome (clean, mixed, heavily contaminated, or rescue). The dashed identity line indicates complete read removal. Rescue barcodes are highlighted by black outlines and represent barcodes retained after design-aware reassignment to the expected genome following removal of conflicting ambient signal. **D,** Total read depth before and after cleaning for the 400 rescue barcodes highlighted in **C**. The dashed line marks the 100-read retention threshold, also required for rescue classification. **E,** Total reads removed by expected well genome, stratified by AmbientMapper call class and colored by the genome to which removed reads mapped.

An important feature of the decontamination process also serves as a diagnostic for technical artifacts, e.g., barcodes whose reads mapped better to a genome other than the one loaded on the corresponding well. To identify them, we compare the dominant genome of every barcode with the genome expected from its plate index. In total, we identified a population of 400 artifacts we denote as “rescue” barcodes (**Fig. 3C**). For each barcode, we defined the expected-genome fraction as the proportion of its confidently assigned reads that map to the genome loaded into its well (*Methods*). Before cleaning, rescue barcodes had a median expected-genome fraction of only 12%. Consequently, after cleaning these barcodes retained few reads (**Fig. 3D**). A pipeline without access to the plate design would either discard these barcodes as genome mismatches or retain them as nuclei assigned to the wrong genome. Plate index aware cleaning instead removes the foreign signal while preserving the barcode, allowing downstream quality control rather than genome mismatch to determine whether the nucleus is retained. Because the ambient pool is dominated by maize, post pooling tagmentation predicts a directional effect, with Arabidopsis wells accumulating maize reads while maize wells remain largely unaffected. Consistent with this expectation, only 5 of 12,442 maize well singlets with at least 100 reads (0.04%) contained a majority of Arabidopsis reads, whereas 679 of 6,701 Arabidopsis well barcodes (10.1%) contained predominantly maize reads (**Table S3**). Together, these results show that modeling ambient contamination in combinatorial indexed data captures its expected directional structure and recovers barcodes that conventional strategies would otherwise discard or misclassify.

A uniform filter would remove a similar fraction of reads from every barcode, whereas removal driven by measured contamination should not. To evaluate this, we counted the reads of each barcode before and after cleaning, keeping its plate index and call class (**Fig. 3C**). Barcodes fell into four groups by the fraction of reads removed. We defined “clean” barcodes as those losing less than 10% of their reads and “heavily contaminated” barcodes as those losing more than half, 45,853 barcodes fell in between, and the 400 rescue barcodes described above lost the most reads in absolute terms, a median of roughly 1,400 each. Removal was also strongly asymmetric, 7.42 million (M) maize reads from Arabidopsis-indexed barcodes against 1.08M Arabidopsis reads from maize-indexed barcodes, 8.5M of 34.14M reads in total (**Fig. 3E**), and the wells losing the most reads were those enriched for barcodes called ambiguous. Together, these results show that AmbientMapper removes reads in proportion to barcode specific contamination rather than applying a uniform filter.

### Robustness Across Diverse Experimental Designs

Maize and *Arabidopsis* diverged approximately 150 million years ago, making their reads relatively straightforward to assign to the corresponding reference genomes. Because AmbientMapper relies on alignment quality, we expected these metrics to lose discriminatory power as reference genomes become more similar, such as among genotypes within a species or recently diverged species. AmbientMapper addresses this limitation at three levels (*Methods*). At the read level, ambiguous reads are retained but down weighted, while reads colocalizing with unambiguously assigned reads receive increased support. At the barcode level, a coherent dominant genome signal reduces support for a spurious doublet call. At the global level, an optional empirical cross mapping profile informs the singlet model of the expected leakage between related genomes, preventing secondary signal from being interpreted as mixing by default. We tested these corrections using three designs of increasing difficulty.

To evaluate these corrections under controlled and increasingly challenging conditions, we first simulated reads from three closely related maize genome assemblies, so that every contaminating read is traceable to its source and the amount of contamination is set by design. Specifically, B73, Il14H and Ki11 assemblies were combined with accessible chromatin regions (ACRs) called from real scATAC-seq B73 data under two distinct conditions. The first condition, which we denoted as “identical-peaks”, represents homologous regions that map one-to-one with Il14H and Ki11 assemblies, and comprise ∼4,000 ACRs with ∼12.9% perfectly conserved across the three genotypes. We also generated a set of “discriminative-peaks”, comprising ∼1,000 ACRs carrying at least one discriminating variant (**Fig. 4A**). Using these two sets of regions as input, we generated synthetic barcodes across 15 contamination regimes spanning empirical contamination α = 0 to 50% for each of two contaminant genomes, setting B73 as the ground truth (**Fig. 4B**). Next, we ran AmbientMapper and compared barcode labels before and after cleaning. As expected, the contaminant detected was proportional to α under both conditions (**Fig. 4C**). Barcode calls also behaved as anticipated in the discriminative-peaks condition, with all true singlets recovered as singlets at α = 0 and a transition to doublet calls from α = 10% onward. In the identical-peaks condition, cross-mapping alone re-classified most uncontaminated singlets into doublet calls (**Fig. 4B**, **Fig. S5E**, **S5F**). Notably, among true singlets called doublet using only the discriminative-peaks, B73 remains the dominant genome in >97% of barcodes through α = 20% (**Fig. S5C**). Importantly, this benchmark also highlights the source of discriminating information. Mapping quality is uninformative among these references and alignment scores result in only marginal separation, leaving edit distance on discriminative peaks as the most informative signal (**Fig. S5I**-**S5K**). For example, when contaminants came from the more closely related Ki11 genome, the identical-peaks condition fails to identify B73 as the top genome across all model configurations. In contrast, discriminative-peaks correctly identifies B73 as the top genome in 27% of doublet calls, and up to 42% across model configurations. With the more distant Il14H, discriminative-peaks retain B73 in about 90% of calls while with identical-peaks B73 falls to 22% (**Fig. S5B, S5C**). These conclusions are insensitive to the model settings, with two exceptions that occur at zero contamination. Enabling the global cross-mapping profile on a panel of only three genomes calls every true singlet as a doublet, while disabling co-localization rescue or up-weighting ambiguous reads results in a 50% reduction in singlet recovery (**Fig. S5**).

**Fig. 4.**
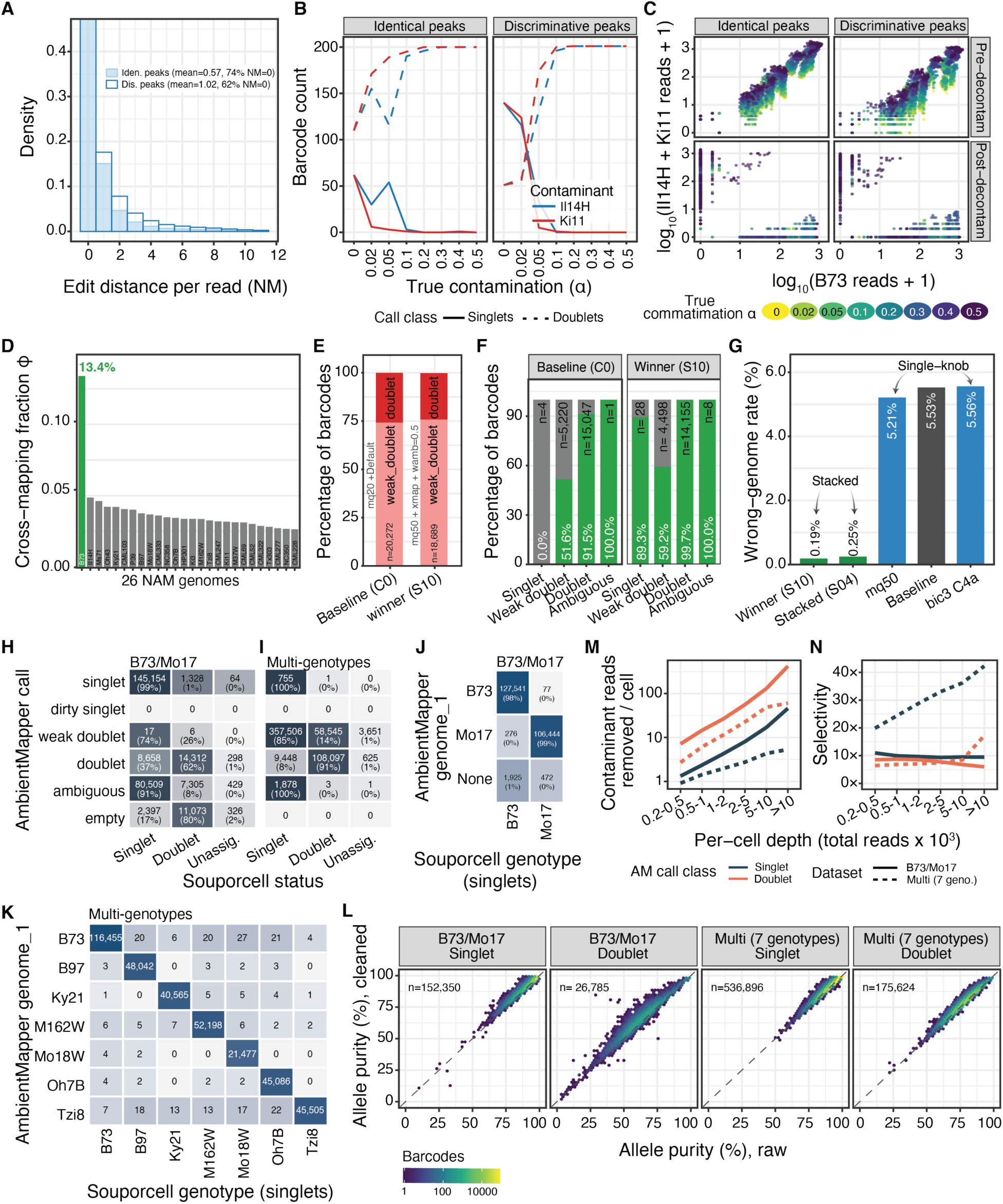
AmbientMapper is robust across diverse experimental designs and genetic backgrounds. **A,** Per-read edit-distance (NM) distributions at zero simulated contamination (α = 0) for two benchmarking tracks: identical peaks (∼4,000 ACRs mappable one-to-one across B73, Il14H, and Ki11; mean NM = 0.57; 74% of reads at NM = 0) and discriminative peaks (“Dis. peak”), containing at least one discriminative variant relative to both Il14H and Ki11 (mean NM = 1.02; 62% at NM = 0). **B,** Numbers of barcodes classified as singlets or doublets across increasing true contamination levels (α), stratified by benchmarking track and contaminant genome (Il14H or Ki11). **C,** Pre-versus post-decontamination read counts for each benchmarking track, colored by true contamination level. **D,** Cross-mapping profile (φ) of B73 reads across the 26 NAM founder genomes. B73 showed the highest mapping fraction (13.4%), exceeding all alternative genomes. **E,** AmbientMapper call composition under the baseline configuration (C0) and the best-performing stacked configuration (S10). Configuration details are provided in Fig. S5 and Fig. S6A. **F,** Dominant genome assignment (genome_1) by AmbientMapper call class for C0 and S10. Because the evaluated library is B73, B73 is the expected dominant genome. **G,** Wrong-genome assignment rate across representative configurations (S10, 0.19%; S04, 0.25%; mq50, 5.21%; C0, 5.53%; C4a, 5.56%). **H,I,** Concordance between AmbientMapper call classes and Souporcell assignments for B73/Mo17 (**H**) and multi-genotype scifi-ATAC (**I**). Percentages indicate the fraction of total cells within each AmbientMapper class assigned to each Souporcell category. **J,K,** Concordance between AmbientMapper top-genome prediction (genome_1) and Souporcell genotype among Souporcell singlets for B73/Mo17 (**J**) and multi-genotype scifi-ATAC (**K**). **L,** Allelic purity before versus after AmbientMapper cleaning. Allelic purity was defined as the fraction of informative reads per barcode that matched the AmbientMapper assigned genome, calculated after WASP correction (p_top1). B73 and Mo17 replicates were pooled. Panels are faceted by AmbientMapper call class. For this analysis, the singlet class combines *singlet*, *dirty singlet*, and *weak doublet* calls (*Methods*). The dashed line indicates y = x. Density above the diagonal indicates increased allelic purity after cleaning. **M,** Number of contaminant reads removed per cell at genotype-informative sites, stratified by AmbientMapper call class and read depth. **N,** Removal selectivity, defined as the contaminant-read removal rate divided by the on-target-read removal rate, stratified by call class, and read depth.

We next asked how the framework behaves when the reference panel is deliberately redundant. We mapped a maize 10x scATAC-seq library derived from the single inbred B73 against 26 high quality maize reference genomes. Because every barcode originates from B73, any non-B73 assignment represents a cross-mapping error. The cross-mapping profile illustrates the difficulty of this setting, with only 13.4% of the reads of B73 barcodes landing on B73 and the rest distributed across other genomes (**Fig. 4D**). Under the selected AmbientMapper model, the wrong-genome rate was reduced to 0.19%, with singlet precision reaching 92.6% (**Fig. 4E to G**, **Fig. S6**). Performance remained strongly dependent on the corrections used to resolve highly redundant references, which we evaluated systematically across parameter combinations and ablations (**Fig. S6**). Importantly, increasing sequencing depth alone did not resolve the problem. Performance without these corrections plateaued at high coverage, whereas the corrected model reached approximately 80% F1 with only 1,000 to 2,000 reads per barcode (**Fig. S6**). Thus, in highly redundant reference panels, cross mapping rather than sequencing depth is the primary limitation.

To test the framework against an orthogonal method on real data, we compared AmbientMapper calls with Souporcell, a single-reference variant-based demultiplexer run in supervised mode, on two multi-genotype scifi-ATAC libraries (**Table S2**), a B73 and Mo17 mixture of 271,876 barcodes and a seven-genotype scifi-ATAC-seq library. Souporcell genotypes barcodes from allele counts at variant sites based on a single reference, while AmbientMapper maps reads competitively against whole genome assemblies, enabling cross-method validation. On barcodes Souporcell calls singlets, the genome assignments match almost perfectly between methods for both datasets (**Fig. 4J**, **4K**). The two methods also agree where each is most confident about class, with singlet barcodes from AmbientMapper being Souporcell singlets in 99% and 100% of cases and confident doublets in 62% and 91% (**Fig. 4H**, **4I**). The residual disagreement concerns class granularity rather than genotype. In the two-genotype mixture, the conservative ambiguous class from AmbientMapper maps predominantly onto Souporcell singlets. In the seven-genotype experiment, the doublet model, calibrated for two components, labels most genetically clean barcodes as weak doublets. These barcodes are still assigned to the correct genotype and are quarantined rather than mis-genotyped, allowing users to determine the most appropriate data retention strategy.

Finally, we tested whether AmbientMapper cleaning improves genotype signal in the multi genotype scifi-ATAC-seq datasets. Because all reads were aligned to a single reference, allele measurements at variant sites are biased toward B73. We therefore estimated per barcode allele purity at the same informative variant sites before and after cleaning, with WASP correction applied to both alignments (*Methods*). Cleaning increased median allele purity across all call classes and datasets, from 97.2% to 97.6% for singlets and from 63.9% to 66.7% for doublets in the pooled B73/Mo17 dataset, and from 95.8% to 96.0% and 80.0% to 80.8%, respectively, in the seven-genotype dataset (**Fig. 4L**). Although the increase in genotype purity was modest, it was consistently positive and supports selective rather than indiscriminate read removal. Consistent with this interpretation, the number of contaminant reads removed per barcode increased with sequencing depth and was highest for doublets (**Fig. 4M**). Across call classes and datasets, contaminant reads were removed at 6 to 42 times the rate of on target reads (**Fig. 4N**).

### Removing ambient contamination restores biological structure

To evaluate the impact of removing contaminating reads on the biological structure of the data, we repeated the cross-species co-embedding of Fig. 1F on the cleaned barcodes post-AmbientMapper. We kept the QC, embedding, and clustering parameters identical, so that any change is attributable to cleaning rather than parameter tuning. We found that the co-projection artifact was removed, with Arabidopsis and maize barcodes resolving discretely (**Fig. 5A, B**). This observation was supported by the Arabidopsis-plate contribution to the embedding, which fell from 34.3% to 3.9% of nuclei (8,540 to 586) post cleaning, and clusters of mixed species composition (fell from all 14 to 3 clusters, **Fig. 5C**). Next, we scored the fraction of each barcode’s nearest neighbors from the other species relative to the expectation under random mixing. Before cleaning, the co-embedding was near random expectation (observed/expected 85-88% across all 36 embedding configurations tested). After cleaning, random mixing fell to 23-27%, a ∼3.5-fold reduction that holds in every embedding configuration (**Fig. 5D**). As expected by the contamination profiles (**Fig. 1**), cleaning removes 75% of the Arabidopsis-plate reads in this design. Thus, to evaluate the effect of the asymmetric cleaning, we next asked whether the surviving data also degraded asymmetrically. TSS fraction was essentially unaffected in maize (median 0.45 to 0.44) and decreased modestly in Arabidopsis (0.47 to 0.42), in both cases remaining well above a traditional threshold (0.2; **Fig. 5E**). The small Arabidopsis decrease is an expected consequence of successful cleaning. Cross-species ambient reads misalign preferentially to conserved TSS-proximal sequence, inflating the pre-clean value. Cleaning leaves FRiP essentially intact, indicating that removed reads largely reflect noise (**Fig. 5F**).

**Fig. 5.**
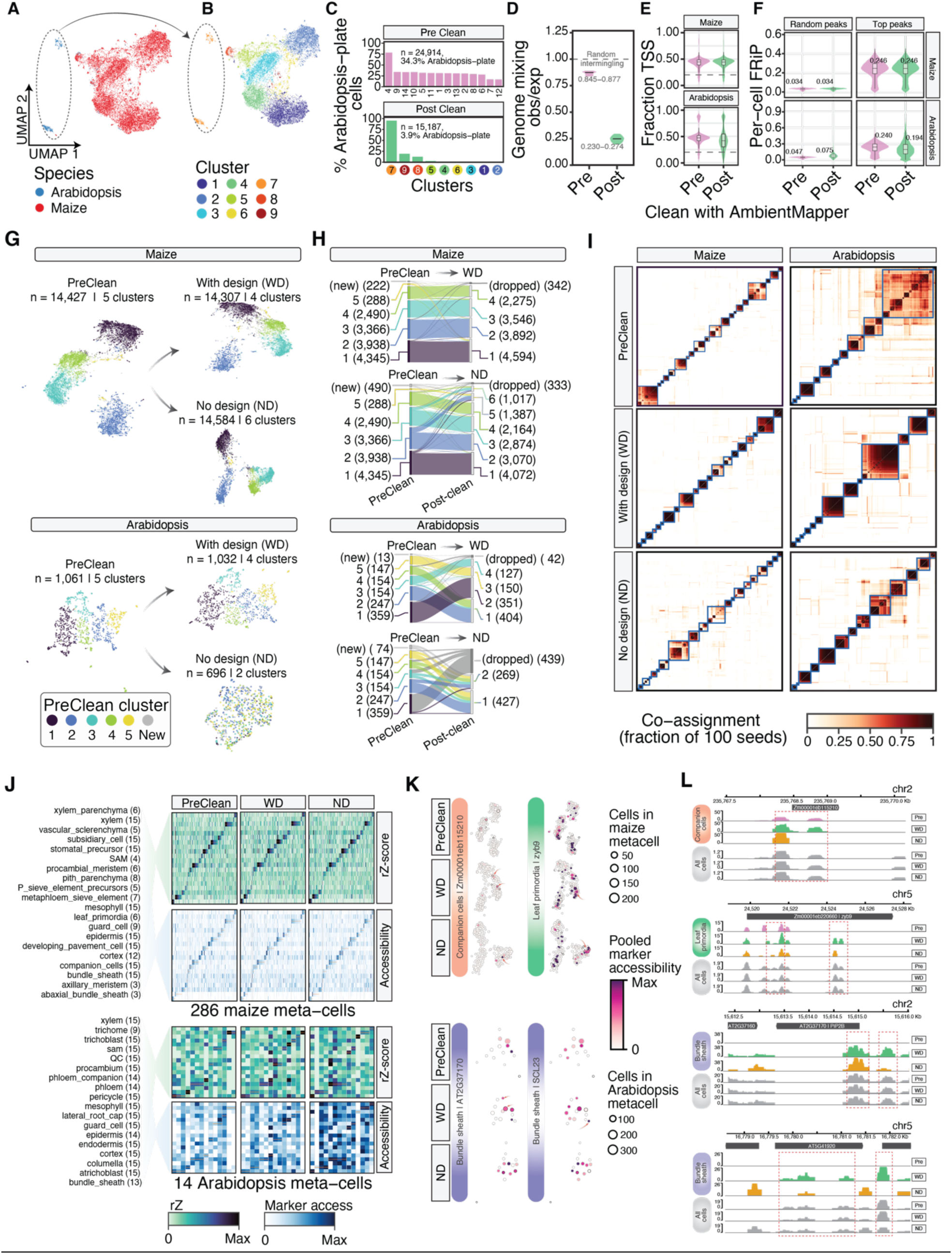
Removal of ambient contamination restores biological structure. **A,B,** Joint embedding of the cleaned datasets using the combined-genome representation (n = 15,187 cells; same configuration as the pre-clean embedding in Fig. 1F), colored by species of origin (**A**) or Leiden cluster (**B**). The dashed outline and arrow indicate the *Arabidopsis* island. **C,** Fraction of *Arabidopsis*-plate nuclei within each cluster before cleaning (top; n = 24,914 cells, 34.3% *Arabidopsis*-plate, 14 clusters) and after cleaning (bottom; n = 15,187, 3.9%, 9 clusters). Clusters are ordered by *Arabidopsis*-plate fraction within each stage. Post-clean cluster colors and labels correspond to **B**. Pre- and post-clean datasets were embedded and clustered independently; cluster identities are therefore not directly paired between stages. **D,** Genome mixing in the joint embedding, quantified per cell as the fraction of nearest neighbors from the other species divided by the expectation under random mixing (Methods). The dashed line indicates observed/expected = 1. Violins summarize 36 embedding-parameter combinations from the grid search (pre-clean, 0.845–0.877; post-clean, 0.230–0.274). **E,** Per-cell TSS fraction for cells retained in both stages (maize, n = 18,591; *Arabidopsis*, n = 8,825; paired). The dashed line indicates the 0.2 QC threshold. **F,** Per-cell FRiP at matched peak number (K = 10,000 peaks per stage), using the cells in **E** and two peak-selection strategies (Methods). **G,** Per-genome UMAPs generated after mapping each library to its corresponding reference genome. Maize (top) and *Arabidopsis* (bottom) are shown before cleaning and after independent well-design-aware (WD) and no-design (ND) cleaning. Cells are colored by their pre-clean cluster of origin; gray indicates unlabeled cells. **H,** Sankey plots showing cluster-membership transitions from pre-clean to WD and from pre-clean to ND for the same objects. Pre-clean clusters are colored as in **G**. **I,** Reproducibility of the meta-cell partition across 100 SEACells runs. Cells are ordered by the consensus partition, with boxes outlining consensus meta-cells. **J,** Heatmaps comparing annotated consensus meta-cells from **I** (columns; 286 maize and 14 *Arabidopsis*) with curated cell-type marker sets (rows; marker number indicated in parentheses) across cleaning stages. Upper blocks show type-level reciprocal z-scores (rZ); lower blocks show pooled marker accessibility rescaled from 0 to 1 within each cell-type row. Orange marks indicate the maximum rZ for each corresponding meta-cell. **K,** Per-genome meta-cell UMAPs, using the embeddings in **G**, displaying representative marker-gene accessibility. Accessibility was capped at the 95th percentile. Black outlines indicate meta-cells assigned to the corresponding marker-defined cell type; arrows highlight meta-cells with increased marker signal after cleaning. **L,** Genome-browser views of the four marker loci shown in **K**. Tn5 insertion coverage is shown as RPM-normalized pseudobulk signal grouped by consensus cell-type assignment within each experimental arm.

We next evaluated cleaning in a per genome analysis, where each plate population was mapped only to its expected reference. We compared cleaning with plate design information, WD, against design free cleaning, ND, to determine whether the experimental design alters the biological signal retained after decontamination. In maize, cleaning had little effect. ACR calls remained relatively stable, with at most a 5.4% reduction relative to precleaning (**Fig. S7A**), and nuclei retained after cleaning showed similar quality profiles to the preclean population (**Fig. S7F**). WD retained 14,307 of 14,427 nuclei, and the preclean cluster structure was largely preserved after cleaning, with a similar pattern under ND (**Fig. 5G, H**). In *Arabidopsis*, where contamination was greatest, the two modes produced markedly different outcomes. WD retained 1,032 of 1,061 nuclei, whereas ND retained only 696 and repartitioned the remaining nuclei into two coarse clusters (**Fig. 5G, H**). Thus, access to the plate design has the greatest impact when ambient contamination is strong enough to obscure the true genome signal.

To assess the impact of cleaning on more fine-scale analysis of cell identity, we repeated the assessment at the meta-cell level, which aggregates highly similar nuclei into a single profile, reducing the effects of sparsity^24^. For each dataset, we ran SEACells with 100 random seeds and built a consensus partition from the co-assignment matrix F, which records the fraction of runs placing each pair of nuclei in the same meta-cell. The meta-cell count was defined at the SEACells-derived value, 286 for maize and 14 for Arabidopsis, and held identical across the three stages, pre-clean, WD and ND, so stage differences cannot arise from differing group counts. Notably, cleaning visibly improves the block structure of matrix F in both species, reflecting greater confidence in meta-cell assignments (**Fig. 5I**). Agreement between runs was scored across all 4,950 seed pairs with the adjusted Rand index (ARI), the chance-corrected proportion of nucleus pairs that two partitions treat concordantly, and the adjusted mutual information (AMI), which applies the same chance correction to their mutual information and is more stable when group sizes are unbalanced^25,26^. Overall, cleaning raised run-to-run agreement sharply in Arabidopsis, from an ARI of 0.47 to 0.72 under WD, with design-free cleaning recovering less structure (0.64). Agreement among maize barcodes exhibited minimal change, from 0.56 to 0.53 ARI, as expected for a majority library with little ambient signal to remove, with AMI performing similarly (**Fig. S8A, B**). The annotation analysis also shows the same asymmetry. The median fraction of seeds supporting each meta-cell’s consensus label rises in Arabidopsis from 0.62 to 0.83 under WD, and the fraction of meta-cells whose consensus label agrees with the modal seed label rises from 71% to 93%, while maize is unchanged at 68% and 82% (**Fig. S8C**). The plate design therefore improves the reproducibility of meta-cell structure, enhancing biological signal, with corrective effects proportional to the extent of contamination.

Finally, we asked whether cleaning recovers interpretable cell-type signal. Each consensus meta-cell was annotated against curated marker sets^27,28^ with a reciprocal z-score and, independently, with pooled per-kilobase marker gene chromatin accessibility. We observed strong agreement between these readouts (Spearman ρ 0.91 in maize, 0.74 to 0.79 in Arabidopsis; **Fig. 5J**). Each meta-cell was labeled with the cell type whose marker set scored highest. Cleaning refined chromatin accessibility at these markers, consistent with the increased co-assignment of nuclei after cleaning. For instance, chromatin accessibility in gene bodies of Arabidopsis bundle-sheath markers AT2G37170 and SCL23 became prevalent in bundle-sheath meta-cells only after the cleaning. Similarly, the maize companion-cell marker Zm00001eb115210 became more specific to companion cells (**Fig. 5K**). Pseudobulk genome browser tracks showcase this recovery at the read level, with improved specificity over the marker gene bodies after cleaning (**Fig. 5L**). All together, these results indicate that cleaning not only removes contaminating reads, but also restores species separation, preserves per-nucleus quality, stabilizes clustering and meta-cell structure, and recovers cell-type-specific biology.

## DISCUSSION

Ambient chromatin contamination in multiplexed single-cell assays is asymmetric, structured, and largely unmodeled by current pipelines. Our two-species plate design made this contamination directly measurable because off-target reads could be identified by genome mapping, while also providing a ground-truth benchmark for same-species pools, where contamination is otherwise hidden. The experiment exposed a limitation of fixed read-count thresholds: at low depth, genuine nuclei can overlap with background-filled barcodes, causing thresholding to either discard real nuclei or retain contaminated barcodes. AmbientMapper addresses this problem by scoring reads competitively across candidate reference genomes and an experiment-derived ambient profile.

A key feature of AmbientMapper is that uncertain reads are classified rather than simply discarded. Read-level classifications and weights are propagated to the barcode, producing graded assignments and enabling contaminating reads to be removed while preserving the underlying nucleus. In contrast, count-based methods such as SoupX^7^, DecontX^19^, CellBender^8^, and CellSweep^21^ infer and remove contamination from the barcode count matrix. These approaches provide corrected matrices for downstream expression or accessibility analyses but do not identify the specific reads or fragments responsible for contamination and therefore cannot directly generate decontaminated coverage tracks or de novo peak calls. Ambimux^20^ models ambient contamination at the read level using donor-informative variants in reads aligned to a common reference, but reports droplet-specific ambient fractions rather than removing contaminating reads from the underlying alignment files. AmbientMapper instead assigns contamination at the read level and propagates those assignments to the barcode, enabling both barcode recovery and reconstruction of cleaned genomic coverage. The distinction is important for shallow nuclei embedded in a large ambient background. Arabidopsis bundle-sheath nuclei provide a clear example. Before cleaning, their relatively small Arabidopsis signal was obscured by maize reads, weakening marker-gene contrast and preventing recognition of the cell type. Removing off-target reads while retaining the barcodes restored both marker contrast and pseudobulk coverage, revealing the expected cell-type-specific pattern (**Fig. 5K**, **5L**). Thus, read-level cleaning can recover low-depth or minority populations while simultaneously restoring the genomic signal associated with those nuclei.

Experimental design provided additional information about both the source and consequences of contamination. Barcode swapping was nearly absent in the reciprocal direction, whereas Arabidopsis-indexed barcodes frequently contained reads mapping unambiguously to maize (**Fig. 1**; **Fig. 3**), supporting ambient chromatin carryover rather than barcode swapping as the dominant source of cross-sample signal. The design also provided a useful prior for decontamination. In well-design-aware (WD) mode, AmbientMapper restricts each barcode to the genome expected from its well, whereas no-design (ND) mode relies only on the genome supported by the reads. Under strong asymmetric contamination, this distinction became important. In the minority pool, ND discarded approximately one-third of the nuclei retained by WD, showing that design-free inference can over-clean populations whose true signal is overwhelmed by ambient chromatin. What determines the size of the ambient pool remains unresolved, although nuclei loading per well and per droplet are plausible contributors that should be tested across extraction and loading conditions.

An unexpected result was the agreement with genotype-based demultiplexing. Although AmbientMapper was not developed as a genotyper, its top-genome assignments reproduced nearly all high-confidence Souporcell singlet calls despite relying on whole-genome competitive mapping rather than variant-level inference. This suggests that, when distinct reference genomes are available, competitive mapping can provide an independent layer of genotype assignment and quality control while simultaneously identifying contaminating reads. More broadly, our results indicate that the measurable cost of ambient contamination is determined by pooling context and is best evaluated within samples before and after read-level cleaning.

Several limitations remain. First, our swapping bound applies to the between-species design and does not quantify well-to-well swapping among samples of the same species, which will require a different experimental design. Second, the current doublet model is calibrated for two components. In a seven-founder pool, some genetically clean nuclei are therefore routed to the weak-doublet class. These nuclei remain assigned to the correct founder but are conservatively quarantined as weak-doublet or dirty-singlet. Class composition should consequently be interpreted with caution in samples containing more than two candidate genomes. Third, although the cleaning effect is consistent across sequencing depth, genotype, and call class, we have not yet modeled these factors jointly.

Finally, AmbientMapper reads standard CB or BC tags without assuming a particular barcode structure. It should therefore be portable to scifi-RNA and other droplet-based assays in which multiple species or sufficiently distinct reference genomes are available. We release scifi-demux alongside AmbientMapper as an entry point for raw scifi-ATAC-seq data, providing plate and well demultiplexing, barcode correction, and adapter handling.

## METHODS

### Plant material

Seeds of the B73 maize inbred were obtained from the USDA ARS-GRIN (https://npgsweb.ars-grin.gov), while seeds of the Arabidopsis ecotype Columbia were sourced from ABRC. Maize plants were grown at 30°C during the day and 20°C at night, under a 16-hour light and 8-hour dark cycle, for 7–8 days until reaching the V2 stage and irrigated with distilled water. Arabidopsis plants were grown at 27°C during the day and 22°C at night, under a 16-hour light and 8-hour dark cycle. For maize, we collected all above-ground organs, while Arabidopsis included all seedlings, including root. All samples were collected in liquid nitrogen and stored for further analysis (−80°C).

### scifi-ATAC-seq library construction

To increase library throughput and the number of technical replicates for each library, we prepared each sample by separating and tagmenting it across 32 independent wells per nuclei extraction, with up to two samples processed per plate. The plate design of every scifi-ATAC-seq library reanalyzed here, including the publicly available libraries, is given in **Table S1**. To construct the scifi-ATAC-seq libraries, we followed previously described scifi-ATAC-seq protocol with a few modifications^4^. Specifically, all material was macerated homogeneously on a mortar and pestle for 4 minutes. Ground material was collected in a pre-chilled 2 mL LoBind® centrifuge tube. We processed immediately to resuspend the ground material in 1 mL of cutter buffer (40 mM of MES-KOH, 40 mM NaCl, 1 M of sucrose, 0.1 mM spermine, 0.5 mM spermidine, 1 mM of DTT, 1% of BSA, and 0.5% of TritonX-100), homogenized by a gentle 5 s (vortex half speed), and incubated on ice for 2 min. The aqueous nuclei slurry was filtered through a 40 μm cell strainer (pluriSelect, cat# 43-10040-40). Then, we pelleted the nuclei by centrifugation (swinging-bucket centrifuge rotor) at 300 rcf for 5 minutes at 4°C, followed by a wash in 500 μL wash buffer (cutter buffer without TritonX-100), and filtered again through a 20 μm cell strainer (pluriSelect, cat# 43-10020-60). We further filtered for chloroplasts, mitochondria, and small debris by passing the nuclei suspension on 1 mL of 35% percoll (wash buffer with 35% percoll) and centrifugation at 500 rcf for 10 minutes at 4°C. Clean pelleted nuclei were resuspended in the 200 μL 2.5x TAPS with detergent buffer (50 mM TAPS-NaOH pH 8.0, 125 mM of MgCl2, 0.1% Tween 20, and 0.01% digitonin).

### scifi-ATAC-seq datasets

To minimize biases arising from genome mapping or software versions, we collected and reanalyzed four datasets representing distinct levels and sources of potential ambient contamination. First, we included a B73 and Mo17 library in which nuclei from both maize genotypes were pooled before processing and treated as a single sample^4^. Second, we analyzed a multigenotype library in which genotypes were processed on the same plate but assigned distinct plate indexes. This design provided independent genotype labels for evaluating between genotype contamination when corresponding reference genomes were available (**Table S2**). Third, we included a library generated from a single maize genotype, which served as a negative control in which cross genotype contamination is not expected (**Table S2**)^27^. Fourth, we generated additional libraries containing B73 and Arabidopsis thaliana, with each species assigned to distinct indexed wells. Increasing the number of wells per species provided technical replication, while the divergent genomes allowed cross species reads to be identified directly. This design therefore provided a positive control for detecting and quantifying ambient contamination.

### Processing of scATAC libraries

Raw paired-end reads of the public maize root scATAC library were demultiplexed with UMI-tools extract^29^, which moves the 16 bp 10x Genomics cell barcode from the index read into the headers of reads 1 and 3. Reads were then aligned independently against each candidate reference genome with BWA-MEM v0.7.17^30^ using the -M flag to mark shorter split hits as secondary. Barcodes observed on fewer than five reads were discarded, and the remainder were corrected against the 10x 737K-cratac-v1 whitelist allowing up to two mismatches and written to the BC tag. Duplicates were removed with Picard MarkDuplicates (broadinstitute.github.io/picard) using BARCODE_TAG=BC and reads with a mapping quality below 10 or carrying multi-mapping evidence were discarded. Surviving alignments were retained as one BAM file per candidate genome.

### Processing of scifi-ATAC-seq libraries with scifi-demux

scifi-ATAC-seq libraries differ from standard single-cell ATAC libraries in carrying two 5 bp Tn5 indexes, one at the start of read 1 and one at the start of read 3, whose combination identifies the well of origin in the plate. We therefore consolidated our previously described preprocessing and post-mapping steps into a single Python package, scifi-demux (github.com/gomezcan/scifi-demux), which adds Tn5 and 10x barcode correction and runs chunk-parallel on either a workstation or an HPC system. It has two modules, step1 for raw reads and step2 for mapped reads.

*step1* takes the 16 bp 10x barcode which is transferred from the index read into the read name with UMI-tools^29^. The two Tn5 indexes are then clipped from the first 5 bases of reads 1 and 3 and appended to the read name with Cutadapt^31^, which in the same pass trims the Tn5 mosaic end (AGATGTGTATAAGAGACAG) from both reads at an error rate of 0.2. Each index is assigned to the closest entry in the plate layout, allowing at most one mismatch per 5 bp index and counting any ambiguous base as a mismatch. Read pairs in which either index exceeds that tolerance are discarded. Corrected reads are written to the sample pool given by the plate index layout, or, when no design is supplied, to one FASTQ pair per plate well.

*Step 2* takes a user supplied mapping plan and aligns the corrected read pairs independently to each candidate reference genome using BWA MEM^30^. Alignments are retained only when reads are properly paired and meet the user defined mapping quality threshold, which was set to 10 in all analyses reported here. The composite barcode is then revalidated. It consists of 26 bases, including a 16 bp 10x barcode followed by two 5 bp Tn5 indexes. Each component is accepted only if it matches its whitelist exactly or can be uniquely corrected by a single mismatch substitution. Reads failing any of these criteria are discarded. Picard MarkDuplicates is then used to remove barcode duplicates. Multimapped reads are additionally removed when MAPQ is below 30 and alternative alignments occur within three mismatches of the primary alignment. Step 2 outputs one BAM file for each candidate genome.

A mapping quality threshold of 10 was applied to every dataset in both routes to ensure that libraries remained comparable. All BWA indexes were built from the NAM founder assemblies^32^ and the TAIR10 Arabidopsis reference^23^. The resulting BAM files were used as input to AmbientMapper. The same alignments were also reduced to single-base Tn5 integration sites in BED format for the quality-control (QC), embedding, and clustering analyses described below. After decontamination, Tn5 integration sites were regenerated from the cleaned BAM files so that analyses before and after cleaning used the same representation.

### Single-cell ATAC-seq quality control and cell calling

Quality control and cell calling was carried out with Socrates^27^. Organellar insertions were removed, and accessible chromatin regions (ACRs) were called per genome with MACS2^33^ using --nomodel --shift -75 --extsize 150 --keep-dup all at a q-value of 0.1. Barcodes were retained as nuclei when they carried at least 200 unique Tn5 integration sites, met library-specific thresholds floored at 0.2 for both the fraction of insertions near TSS and the fraction within ACRs, held an organellar fraction below 0.2. Counts were summarized as a binary barcode-by-feature matrix over 500 bp windows. Called ACRs were used for quality metrics only and not as embedding features.

### Quantification of ambient contamination

Contamination in the interspecies library was computed from AmbientMapper’s per-barcode winner-read counts. A barcode’s off-target fraction is the fraction of its winner reads assigned to a genome other than the one its plate position expects, summarized per plate over barcodes with more than 10 winner reads. The dominance ratio was rebuilt from the integer winner counts as (k1 + 1) / (k2 + 1) on the best and second-best genome, rather than taken from AmbientMapper’s model-estimated ratio, which is undefined for barcodes with no second-genome read and would drop exactly the shallow barcodes the panel exists to show. To exclude mapping strategy as the cause of the asymmetry, the same library was re-analyzed on a concatenated maize and Arabidopsis index, assigning reads to species by chromosome, and per-species quality metrics on that object were compared with two-sided Wilcoxon rank-sum tests for location shifts and Kolmogorov-Smirnov tests for distributional differences.

### Dimensionality reduction, clustering, and cross-species co-embedding

The binary window-by-nucleus matrix was normalized by term frequency-inverse document frequency (TF-IDF, scale factor 10,000), reduced by singular value decomposition (SVD), embedded with UMAP^34^ and clustered with the Leiden algorithm^35^ on the SVD embedding using Seurat^36^. Every embedding reported here used 20 SVD components, 30 nearest neighbors, a minimum distance of 0.3, a minimum cluster size of 50 nuclei and a resolution of 0.5. That configuration was held fixed for comparability rather than chosen by optimization. A grid of 45 configurations spanning SVD depth, neighborhood size and minimum distance was scored on the concatenated-reference object by an explicit combination of neighborhood preservation, genome mixing, and correlation between embedding coordinates and quality covariates. Species remained intermingled across the whole grid, so a single configuration was applied to every object rather than re-optimizing each, ensuring that a difference between panels reflects the variable under study rather than a change in tuning.

Cross-species co-embeddings were built from the two per-genome 500 bp window matrices by stacking them over the union of barcodes, prefixing features by genome and zero-filling features absent from a nucleus’s genome, so that every nucleus occupies one shared feature space without competitive read assignment. Pre-cleaning and post-cleaning objects were processed identically, the post-cleaning object from WD-cleaned reads. Species intermixing was quantified per nucleus as the fraction of its 15 nearest neighbors carrying the other species label, averaged over nuclei. Because that raw fraction is confounded by species composition, which cleaning itself changes, it was divided by its expectation under random intermingling, 2p(1 - p) at minority fraction p, and this observed over expected ratio is reported throughout. For the robustness analysis it was recomputed under all 36 configurations of the co-embedding grid. Per-nucleus TSS fraction and peak-matched FRiP before and after cleaning (**Fig. 5E, F**) were compared on these objects, reducing the analysis to only nuclei present in both stages (18,591 maize and 8,825 Arabidopsis). Peak-matched FRiP was computed at a common 10,000 peaks per stage under two selection rules with opposite biases, a uniform random draw, and the top peaks by MACS2 score, together providing upper and lower bounds on the true effect and both are reported.

### Synthetic read-level benchmark

To measure classification and decontamination where truth is exact, we built a benchmark in which the genome of origin and the contamination status of every read are known. Accessible chromatin regions called from the public maize root library’s B73 alignment were mapped onto the Il14H and Ki11 NAM founder assemblies^32^ with minimap2^37^, keeping peaks with orthologous placements in all three genomes. Two marker tracks were defined on that shared universe, one using every orthologous peak and one restricted to peaks where both contaminant genomes are expected to carry at least one discriminating variant per read. Paired-end 75 bp reads were simulated from each genome’s peak sequences with ART^38^ and assembled into 208 template barcodes, comprising 150 singlets spanning six depth bins from 250 to 7,500 reads, 48 doublets and 10 empty barcodes. From that fixed template set, 15 datasets were generated per track, one uncontaminated baseline and seven contamination levels from α = 0.02 to 0.50 for each of the two contaminant genomes. Cell reads are identical across the 15 datasets and only contaminating reads differ, so each dataset carries a per-read truth table. All synthetic datasets were processed through the same multi-reference pipeline as the real libraries and evaluated against truth at the read and barcode level.

### AmbientMapper configurations evaluated

Every analysis uses the canonical configuration C0 unless a named variant is stated. Variants are single or stacked changes to nine parameters, the genotyping MAPQ floor, the XA cap, top-K reclassification, the winner-mass discount, co-localization rescue, the cross-mapping profile φ, the number of ambient-profile refinement iterations, the ambiguous-read weight and the BIC margin. The setting of every parameter in every configuration is tabulated in **Fig. S5A** for the synthetic benchmark and **Fig. S6A** for the 26-genome stress test.

### Cross-mapping stress test on a 26-genome reference panel

As a stress test of cross-mapping behavior we used a published scATAC-seq library from a single maize inbred ^27^, in which every barcode is a B73 singlet by construction, and mapped it independently against all 26 NAM founder assemblies^32^ before processing it through the same AmbientMapper pipeline. Any barcode whose assigned genome is not B73 is therefore a cross-mapping error. We report the wrong-genome rate, the fraction of retained barcodes whose top genome is not B73, together with singlet precision and F1 stratified by per-barcode depth, each computed over the barcodes surviving that configuration’s own read filters. The configuration factorial was run on a depth-balanced subsample of 996 barcodes, and the configurations of interest were then evaluated at full library scale.

### Independent genotype assignment with Souporcell

As an orthogonal, variant-based reference we ran Souporcell (version v2.1)^14^ in supervised mode (using known variant call) on the two-genotype and the seven-genotype libraries, with k set to the number of pooled genotypes. Only barcodes with at least 500 fragments were assayed. Souporcell clusters were matched to reference SNPs by Pearson correlation between cluster and corresponding references. B73 is the reference assembly and is homozygous reference at essentially every panel site, so correlation was uninformative for it. Thus, the B73 label was instead matched to the unassigned cluster with the highest homozygous-reference fraction. A single remaining genotype was assigned to the single remaining cluster by elimination. Souporcell and AmbientMapper calls were compared in two ways, the projection of AmbientMapper call classes onto Souporcell singlet, doublet and unassigned status, and genome agreement restricted to Souporcell singlets. Because Souporcell genotypes barcodes from allele counts at variant sites whereas AmbientMapper maps reads against whole-genome references, the two share no genotyping machinery and their agreement is independent validation.

### Allele-level validation of decontamination

To measure the effect of cleaning at allele resolution without reference-mapping bias, raw and cleaned BAM files were corrected with WASP^39^. WASP was run in paired-end mode on coordinate-sorted alignments. Allele counts at informative sites were then counted identically on the corrected raw and cleaned alignments. The same barcodes and sites appear on both sides of every comparison. Per-barcode allele purity was estimated as the fraction of a barcode’s allele-informative reads supporting the genome AmbientMapper assigned, with barcodes stratified by dataset, call class and per-barcode depth in six bins from 200 reads to more than 10,000. Weak doublets were pooled with singlets, because at informative sites the genome pair assigned to a weak doublet is dominated by one genome rather than split evenly (**Table S4**). This pooling affects 0.02% of the B73 and Mo17 singlet class and 99.8% of the multi-genotype singlet class, so the two singlet groups are not comparable in composition. Finally, contaminant reads removed per cell is the mean number of contaminant-allele reads cleaning removed per barcode within each stratum. Removal selectivity is the contaminant-allele removal rate divided by the on-target removal rate at the same sites. Values close to 1 indicate removal blind to origin.

### Cleaning modes and the per-genome analysis

AmbientMapper decontamination was applied in two modes to the same input. Under WD the plate design sets each barcode’s expected genome, and under ND no design is used and reads consistent with the barcode’s top-ranked genome are retained. The two are independent treatments of the same raw data, not sequential steps. For the per-genome analyses (**Fig. 5G** to **L** and **Fig. S7**) each plate library was demultiplexed by its plate index, mapped to its own reference alone, maize B73 or Arabidopsis TAIR10, and passed through the quality control described above. Thus, we generated three analyses per genome, PreClean, WD, and ND. The embedding and clustering configuration was selected separately for each genome on a stability and purity scan, rather than held fixed as in the co-embedding, and was then applied unchanged to that genome’s three analyses. Per-nucleus TSS fraction was compared across the three stages. Per-nucleus FRiP was recomputed on the nuclei present in all three stages, 18,253 in maize and 1,268 in Arabidopsis, subsampled at random to 10,000 for maize. Peak-matched FRiP (**Fig. S7D**) was computed at the smallest of the three stage peak counts under the same two selection rules. In the cross-stage cluster comparisons (**Fig. 5G** and **H**), each barcode was labeled per PreClean cluster and reported membership flows from PreClean to WD and to ND separately. Post-cleaning labels come from independent Leiden runs and are not identified across stages. Barcodes present in only one stage are reported as dropped or new. The stage analyses (PreClean, WD, and ND) are not nested, because cleaning both removes barcodes and rescues nuclei independently.

### Consensus meta-cells

Meta-cells were computed with SEACells v0.3.3^24^. The kernel was built on the object’s SVD embedding, frozen from the clustering step (10 waypoint eigenvectors and 15 neighbors), so the accessibility matrix only aggregates counts. Two confounds were fixed by design. The number of meta-cells was pinned to the value derived on the PreClean, 286 for maize and 14 for Arabidopsis, and held identical across the analyses. Aggregated fragment counts were then downsampled to a common budget set by the shallowest stage of each genome (14,235 fragments for Arabidopsis and 44,658 for maize). Each object was run with 100 random seeds. The co-assignment matrix F, was summarized into one consensus partition by average-linkage hierarchical clustering on 1 minus F, cut at the pinned meta-cell number. A consensus was accepted only if its mean adjusted Rand index (ARI) to the seed partitions exceeded the mean seed-to-seed ARI. Run-to-run reproducibility was quantified as the mean ARI over all 4,950 seed pairs, with adjusted mutual information as a robustness check. Confidence intervals for contrasts were estimated by percentile bootstrapping on resampling runs with 10,000 iterations. For each consensus meta-cell, label support was recorded as the fraction of seeds supporting its type label.

Annotations were performed based on marker-level accessibility, as previously done for barcode clusters^27^. Briefly, gene-level accessibility was computed by counting Tn5 insertions over gene bodies extended by 500 bp on each side and normalizing per kilobase of the gene model. Meta-cell profiles were built by aggregating that raw per-kilobase matrix over the consensus meta-cells and then scaling to counts per million. Marker sets combined curated maize and Arabidopsis markers^27,40^, restricted to the 15 most informative markers per cell type. Each gene by meta-cell pair received a reciprocal z-score combining the gene’s z-score across meta-cells with the meta-cell’s z-score across genes, multiplied as the square root of the product of their positive parts. Each meta-cell’s provisional label is the cell type with maximal aggregate score, weighted by seed support. For browser views, single-base Tn5 insertion sites were extended by 100 bp on each side and pooled by consensus type call within each meta-cell with BEDTools *slop*^41^, converted to reads-per-million coverage with BEDTools *genomecov* written as BigWig with UCSC wigToBigWig^42^ and rendered with pyGenomeTracks^43^.

### Barcode genotyping with multi-reference mapping with AmbientMapper

Profiling large numbers of nuclei is a central advantage of pooled-genotype and combinatorial indexed single-cell ATAC-seq. AmbientMapper genotypes barcodes from reads mapped independently against several reference genomes, as an alternative to SNP-based assignment, and uses the same evidence to remove contaminating reads. It runs as four steps that can be executed independently or as one fully parallel job, locally or on an HPC system.

*Step 1, extract.* Multi-reference BAM files are reduced to per-genome tables holding, for each read and barcode, the alignment score (AS), mapping quality (MAPQ), mismatch count (NM) and number of alternative hits (XAcount). Reads are retained if they carry a valid barcode flag, MAPQ of at least 10 and XAcount of at most 2, all user-tunable. Input BAM files must come from BWA-MEM^30^. The genotyping step later applies its own stricter thresholds, given in the configuration anchor below

*Step 2 filte.* Each per-genome table is reduced to primary alignments, deduplicated by read and barcode, and restricted to barcodes named in the sample design. Tables are then split into user-defined barcode chunks so that assignment runs in parallel.

*Step 3 assign.* AmbientMapper decides, for each read, whether one genome clearly outscores all others. Empirical score distributions are learned from the data itself in a streaming four-pass algorithm that scans each per-genome table at most once per pass, which bounds memory at experiment scale. Throughout, candidate alignments are ranked by fewer mismatches first, then higher alignment score, then higher mapping quality.

*Pass A* learns global decile boundaries of winner quality for AS and MAPQ.

*Pass B* learns, within each decile, the empirical distribution of the gap between the best and second-best alignment.

*Pass C* scores every read against learned distributions and labels it a winner when only one genome aligns it, when the runner-up carries strictly more mismatches, or when either tail probability falls at or below α (default 0.05). Reads failing all three tests are labeled ambiguous. Every read nonetheless carries a putative winner, the top-ranked candidate genome, so the ambiguous label records the absence of statistical separation rather than the absence of a preferred genome.

*Pass D* then rescues ambiguous reads by spatial co-localization. Even between highly similar references, individual reads carry distinguishing differences. Consequently, the per-read winner is noisy but on average unbiased toward the true source genome. Confident winners from one barcode therefore cluster within the loci that barcode occupies, and an ambiguous read from the same barcode falling inside such a cluster very likely shares their origin. AmbientMapper groups reads by barcode, genome, and chromosome, indexes the fragment endpoints of confident winners, and promotes an ambiguous read to rescued when either of its endpoints falls within 500 bp (configurable) of a winner endpoint in the same barcode and genome. Both endpoints are tested independently, so that paired-end fragments anchored by only one mate are captured. The class space at the end of Step 3 is therefore winner, rescued and ambiguous. Disabling rescue (configuration C1c) is evaluated on both the synthetic benchmark and the 26-genome stress test (**Fig. S5**, **Fig. S6**).

*Step 4 genotyping.* Three choices distinguish this stage from prior ambient-removal tools. First, cross-mapping reads are kept with a class flag rather than discarded, as in Xenome^44^ and Disambiguate^45^, or redistributed across candidates by expectation maximization, as in PathoScope^46^. Every read enters the barcode model with a definite genome and a confidence annotation, so uncertain evidence can be discounted without losing its direction. Second, cross-mapping is corrected at three independent scales, a class-keyed reliability discount at the read level, a data-driven winner-mass discount at the barcode level, and an optional global cross-mapping profile learned from the experiment’s own confident singlets. The three address distinct failure modes and can be engaged separately. Third, the doublet mixing ratio ρ is kept independent of the ambient fraction α rather than absorbed into it. Ambient-removal-only tools such as CellBender^8^ collapse doublets into high-α singlets, because reads attributable to a secondary genome have nowhere else to go in the model. Carrying α and ρ separately costs one degree of freedom and makes doublet recovery explicit, which suits pooling designs where the doublet rate is itself a primary readout.

### Barcode-level model selection

AmbientMapper aggregates read-level evidence into a per-barcode expected count for each genome and compares three generative models under an explicit ambient component. The empty model treats all reads as ambient. The singlet model assigns the barcode to one genome with an ambient fraction α. The doublet model assigns it to two genomes mixed in ratio ρ, again with ambient fraction α. Reads classified as ambiguous enter these likelihoods multiplied by a reliability weight *w_ambiguous*, set to 0.1 in every run reported here (default 1.0), against *w_confident* = 1.0 for winner and rescued reads. Because the discount multiplies the ambient mass as well, it changes only how much a read contributes, not which genome it prefers. The ambient profile η is estimated from the barcodes of lowest effective depth. An optional refinement by Jensen-Shannon divergence to the current η is available. Model evaluation is restricted to each barcode’s top K genomes by expected count, where K is 3 by default and was set to 2 here (see configuration anchor). Genomes outside that set still contribute to η but cannot themselves be called.

Fitting α over 0 to 0.5 in steps of 0.02 and ρ over 0.1 to 0.9 in steps of 0.05, models are compared by the Bayesian Information Criterion, BIC = −2 log *ℒ* + *k* log(*n*b), with *k* = 0 for the empty model, 1 for the singlet and 2 for the doublet. Genome identities are treated as model structure rather than as free parameters. Immediately before scoring, a barcode-local winner-mass discount scales every non-dominant candidate genome by its share of confidently assigned mass relative to the dominant genome. A genuine second component keeps most of its mass and remains a viable doublet partner, whereas a candidate supported only by cross-mapped ambiguous reads is compressed toward zero, and the doublet model loses the evidence base it would have used to outscore the singlet. Removing this discount together with top-K re-classification and co-localization rescue (configuration C1g) collapses the doublet class, routes more barcodes to ambiguous than any other configuration and drops singlet precision to 47.9% in the 26-genome stress test (**Fig. S6**).

### Two-stage decision and call vocabulary

Barcodes with fewer than *min_reads* unique reads (C0 value 5) are labeled “low reads” and fit only under the empty model. The remainder pass two decisions. A barcode is called empty only when it satisfies both a BIC separation of at least *empty_bic_margin* (C0 value 6) and structural diffuseness, meaning *p*_top1_ at most *empty_top1_max* (C0 value 0.6) and a top1 to top2 ratio at most *empty_ratio12_max* (C0 value 2). An optional third term comparing the barcode composition to η is available but was not enabled for any analysis reported here. The conjunction is deliberate, because a low-depth barcode concentrating its reads on one genome is structurally unlike ambient and is protected from the empty call, which in turn protects rare cell types that happen to be shallow. For barcodes passing that gate, the singlet and doublet models are separated by *bic_margin* (C0 value 6). A difference below that margin returns an “ambiguous” core call, with the better-scoring model reported and a near-tie flag set when the gap is at most *near_tie_margin* (C0 value 2). A singlet is refined to “singlet” when *p*_top1_ is at least *single_mass_min* (C0 value 0.6), the top1 to top2 ratio is at least *ratio_top1_top2_min* (C0 value 1.5) and no near-tie flag is set, and to “dirty singlet” otherwise. A doublet is refined to doublet when the read-mass share of its minor genome, min(p_top1, p_top2), reaches doublet_minor_min (C0 value 0.20, equal to the default), and to weak doublet otherwise. The call space is therefore “empty”, “singlet”, “dirty singlet”, “doublet”, “weak doublet”, “ambiguous” and “low reads”.

### Optional mechanisms for high-redundancy reference panels

Four switches act on the read evidence before model selection. Each is given with its library default and the value used here.

*Top-K read re-classification.* When enabled, the assignment labels are read a second time with each barcode restricted to its top K genomes by expected count, and every read is re-labeled by a three-tier rule. A read keeps its winner label when exactly one candidate carries it, is promoted to winner when exactly one candidate carries the rescued label, and otherwise goes to the candidate with the unique highest composite score, AS + MAPQ − NM with unit weights, staying ambiguous on a tie. Expected counts are rebuilt from the re-labeled reads, whereas η and, if present, φ keep their full-panel estimates. The pass is off in the library and K defaults to 3. It was on with K = 2 in every run reported here. Its joint ablation with the winner-mass discount and co-localization rescue is configuration C1g (**Fig. S6**).

*Cross-mapping profile φ.* For each anchor genome g1, barcodes whose top genome is g1, with at least 500 unique reads and a top1 to top2 ratio of at least 2, form a confident singlet set. Over that set, the mean expected count on g1 and, for every other genome, the mean excess of expected count over unique reads are column-normalized into φ(g | g1), the fraction of a g1 singlet’s mass landing on g. When the largest off-diagonal entry χ(g1) exceeds 0.05, the singlet likelihood for g1 becomes Σg φ(g | g1)·Lg(r), so mass on relatives is anticipated rather than read as mixing. The doublet likelihood is unchanged. The three thresholds are library defaults, unchanged here. The library estimates φ by default. We disabled it (--no-xmap) in C0 and in every dataset-level run and enabled it only in the configurations marked **Fig. S5A** and **Fig. S6A**, among them S04 and S10 (**Fig. 4D-4G**).

*Winner-only posteriors.* Each read can instead be collapsed to its single best candidate by composite score, which receives unit mass against an ambient pseudocount ε (library default 0.001). This flattens minor components and biases η. The library enables it by default. Every run reported here used --no-winner-only.

*Promiscuous-read filter.* When --max-hits and --hits-delta-mapq are both set, an ambiguous read with more than the capped number of genomes within δ MAPQ of its best hit is discarded before posteriors are computed. Winner and rescued reads are never affected. Both are unset by default and were never set here.

### Configuration anchor

Unless stated otherwise, every analysis in this manuscript uses the canonical configuration C0, defined by a genotyping MAPQ floor of 20, an XA cap of 0, top-K re-classification enabled with *K* = 2, the winner-mass discount enabled, co-localization rescue enabled with a 500 bp window, the cross-mapping profile disabled, no ambient-profile refinement (see Barcode-level model selection), reliability weights *w_ambiguous* = 0.1 and *w_confident* = 1.0, winner-only posteriors disabled, a softmax temperature β = 10 (library default 1), *bic_margin* = 6, *empty_bic_margin* = 6, an α grid of 0.02 over 0 to 0.5, and a ρ grid of 0.05 over 0.1 to 0.9. The cross-mapping profile φ was disabled in C0 and in every dataset-level run, and is enabled only as indicated **Fig. S5A** and **Fig. S6A**. Robustness to systematic perturbation of these settings, including ablation of co-localization rescue (C1c), joint ablation of the winner-mass discount, top-K re-classification, and co-localization rescue (C1g, 26-genome test only), *w_ambiguous* between 0.05 and 0.5, and *bic_margin* between 3 and 6, was evaluated across the three-genome synthetic benchmark with informative and non-informative marker tracks (**Fig. S5**) and the 26-genome stress test on a B73-only library (**Fig. S6**).

### Decontamination and BAM purification

After per-barcode genotyping, AmbientMapper performs a model-informed read-level cleaning step (**decontam**) and then applies its output to alignments (**clean-bams**). Decontamination is deliberately decoupled from the genotyping internals, so read-level decisions depend only on the per-barcode call and on assignment evidence already computed in Step 3. decontam writes a per-read drop list, a per-barcode policy table, a post-clean barcode table, an augmented calls table used for all downstream analyses, and pre-clean and post-clean summaries of per-barcode genome composition that quantify cross-genome mixing without a combined reference.

*Allowed sets.* Each barcode receives an *AllowedSet* of genomes whose reads may enter the cleaned BAM, together with an action, “keep cleaned” or “drop barcode”. Barcodes called empty receive an “empty” *AllowedSet* and are dropped. When the plate design is used (WD), the expected genome comes from the plate design, matched through a barcode key taken from a configurable segment of the barcode string (--design-bc-mode, --design-bc-n), which may be the part before the first dash, the whole string, or its first or last n characters. Here we used the final 10 bases (--design-bc-mode last, --design-bc-n 10), which carry the Tn5 well index as a 5-base column barcode followed by a 5-base row barcode. Strict design enforcement was used throughout, so a barcode whose key is absent from the design map is dropped whatever its call. When the plate design is not used (ND), there is no expectation and the AllowedSet is built from the call itself.

The AmbientMapper defaults retain both genomes of a mixture, *top12* for doublets and for “ambiguous” barcodes, and rescue ambiguous barcodes to the design-expected genome, “design rescue”. We took a more conservative approach, using top1 for doublets so that a barcode called as a doublet contributes only one genome, retaining design rescue for ambiguous barcodes, and keeping the default top1 setting for weak doublets. The ND runs are identical except that ambiguous barcodes follow “top1 rescue”. Every retained barcode therefore carries a single-genome *AllowedSet*, the design-expected genome under WD and the top-supported genome g₁ under ND. Barcodes called “low reads” carry no genotype, so under ND they are dropped at this stage, while under WD they inherit the expected genome and are resolved by the retention gate below.

*Read-level rule.* A read is eligible for decontamination when the tail probability of its winner margin is ≤ decontam_alpha, set to 0.05 in all analyses reported here, or when this probability is undefined because no competing genome produced a comparable alignment. Among eligible reads, a read is added to the drop list only when its winner genome falls outside the AllowedSet assigned to its barcode. Reads classified as ambiguous or rescued, together with winner reads that do not meet the confidence criterion, are retained. Cleaning is therefore restricted to reads whose genome assignment is sufficiently confident and inconsistent with the barcode assignment. Preclean and postclean composition tables are calculated using the same set of eligible reads. An optional safe keep rule, safe_keep_delta_as, retains an otherwise removable read when its best alignment to a genome within the AllowedSet is within a specified alignment score difference of its best alignment outside the AllowedSet. This option is disabled by default and was set to 3 here, where it retained no reads.

*Barcode retention.* After read-level filtering, each barcode is re-evaluated on its surviving allowed evidence against two criteria, a minimum number of allowed reads (min_reads_post_clean, default 100) and a minimum allowed fraction (min_allowed_frac_post_clean, default 0.90). Barcodes failing these criteria are recorded for dropping. This gate is separate from the genotyping-stage gates and is where design enforcement takes effect. A barcode whose reads are dominated by a genome other than the one the design expects loses that evidence to AllowedSet filtering, falls below the read minimum, and is removed here rather than at genotyping. In the runs reported here the read minimum was the binding criterion. Because safe keep spared no reads, every read that survived filtering carried its winner genome inside the AllowedSet. Thus, the allowed fraction was 1 for every retained barcode and the fraction criterion removed no barcodes.

### BAM filtering

Finally, the drop list is applied with **clean-bams**, which removes alignments whose read query name appears in the list. Two properties follow. The same drop list applies identically to any BAM built from the same FASTQ files, including the same reads mapped to several references, and the result does not depend on aligner version, reference assembly or coordinate system. Because cleaning acts on reads rather than on a cell-by-feature matrix, downstream quality metrics such as transcription start site enrichment, fraction of reads in peaks and fragment size distribution are computed directly from the cleaned alignments rather than reconstructed from a denoised count matrix.

## DATA AVAILABILITY

Previously published data reanalyzed in this study are available from the NCBI Sequence Read Archive. The scifi-ATAC-seq libraries^4^ are under BioProject PRJNA996051, runs SRR25320545 to SRR25320547 and SRR25320539 to SRR25320541 for the two B73/Mo17 replicates and SRR25320542 to SRR25320544 for the multi-genotype library. The maize B73 (root) scATAC-seq library^27^ is under BioProject PRJNA648930, sample GSM4696884, of which the sequencing run SRR12331466 was reanalyzed here. The maize and Arabidopsis scifi-ATAC-seq library generated in this study has been deposited under accession [XXX]. A full dataset inventory with accessions is given in **Table S2**.

## CODE AVAILABILITY

AmbientMapper is an open-source Python package available at https://github.com/gomezcan/ambientmapper (v0.1.0, MIT license). scifi-demux, the preprocessing package for scifi-ATAC-seq libraries, is available at https://github.com/gomezcan/scifi-demux (v0.1.3, MIT license). All workflow scripts, run configurations and figure-generating code for the analyses in this study are available at https://github.com/gomezcan/ambientmapper_manuscript.

## USE OF GENERATIVE AI

The authors wrote all the main text. Anthropic and OpenAI were used to edit the manuscript for clarity and to review and document code. All AI-assisted output was reviewed and approved by the authors, who take full responsibility for the content.

## ACKNOWLEDGEMENTS

We wish to thank the Advanced Research Computing at the University of Michigan, Ann Arbor for providing computational resources and technical support. This work was supported by the National Institutes of Health (R00GM144742) and the Office of the Vice President for Research at the University of Michigan (A.P.M.). F.G.C. was supported by the Momental Foundation and is currently funded by the Michigan Pioneer Fellows Program.

## CONTRIBUTIONS

A.P.M. conceived the study. F.G.C. performed experiments. A.P.M. and J.D.W. designed and supervised analysis. F.G.C., L.J., and A.P.M. analyzed the data. F.G.C. and A.P.M. wrote the manuscript with input from all authors. All authors approved the final manuscript.

## Supplemental Figures

**Fig. S1.**
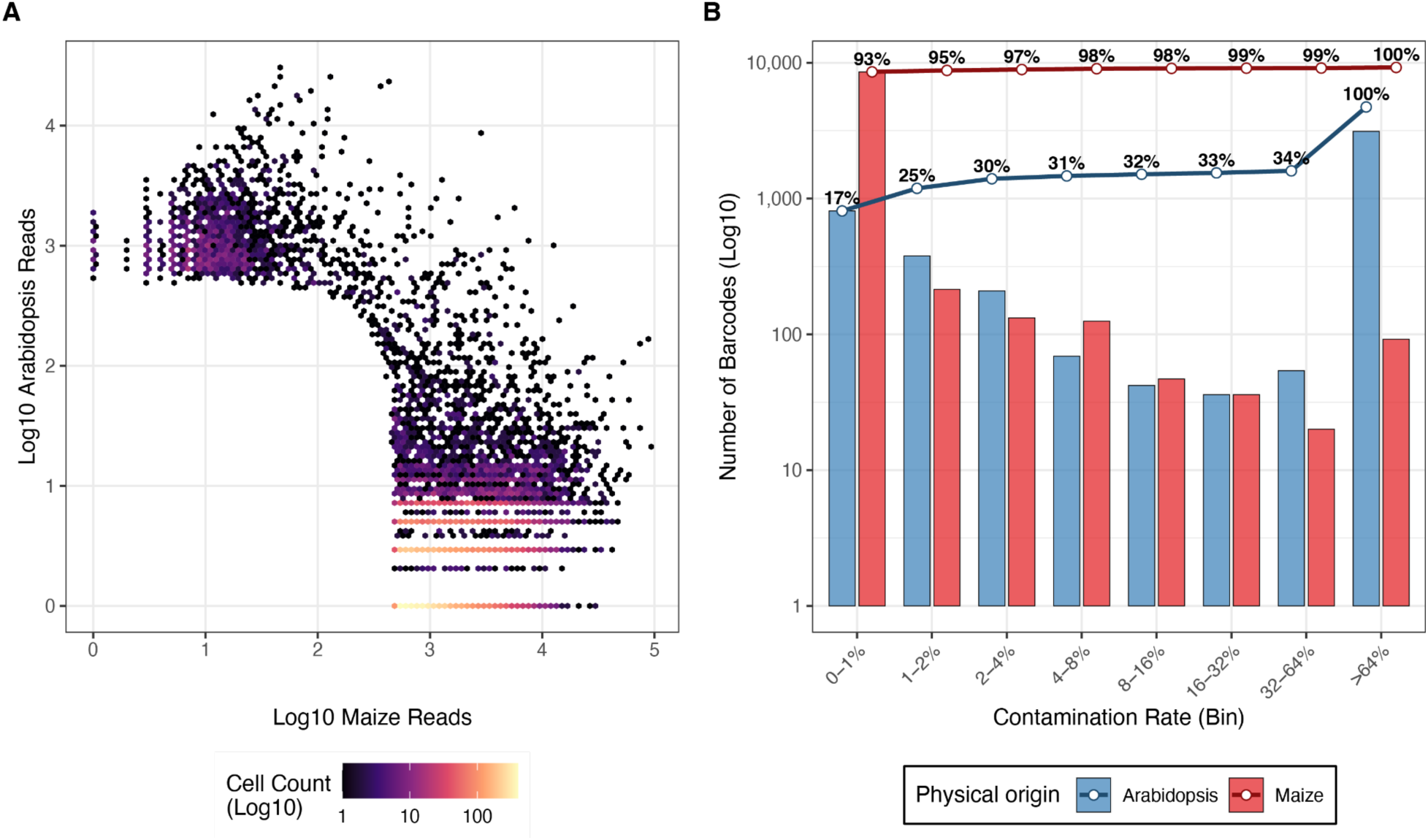
Cross species ambient contamination is robust to mapping strategy. Raw reads were independently aligned to a concatenated *Zea mays* and *Arabidopsis thaliana* reference to test whether the contamination observed in Fig. 1 depended on the competitive winner read strategy. **A,** Barnyard density plot showing unique reads mapping to maize versus *Arabidopsis* for each barcode. Mixed species profiles persisted under concatenated reference mapping, indicating that the observed contamination is not specific to the alignment strategy. **B,** Distribution of per barcode contamination fractions, stratified by physical plate origin. Bars show barcode counts across contamination bins, and lines show cumulative percentages. *Arabidopsis* indexed barcodes contained substantially higher fractions of maize derived reads, recapitulating the asymmetric contamination observed in the primary analysis.

**Fig. S2.**
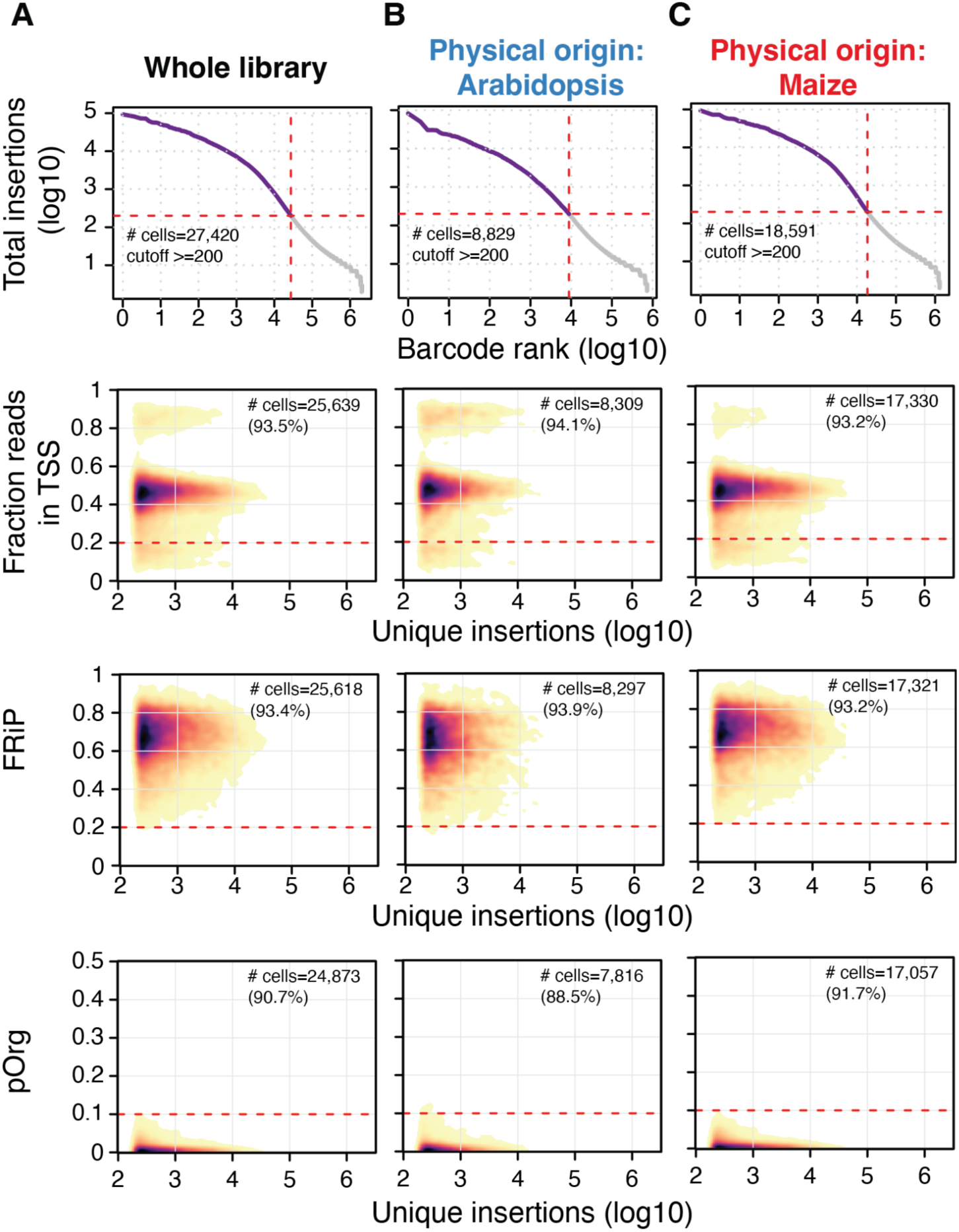
Standard quality control metrics reveal systematic bias against Arabidopsis cells under ambient contamination. Standard cell calling and quality control were applied to reads aligned to the concatenated *Zea mays* and *Arabidopsis thaliana* reference. **A to C,** Results for the full library (**A**), barcodes indexed in *Arabidopsis* wells (**B**), and barcodes indexed in maize wells (**C**). Top row shows barcode rank (“knee”) plots based on unique Tn5 insertions. Lower rows show sequencing depth versus the fraction of reads at transcription start sites (TSS), the fraction of reads in accessible chromatin regions (FRiP), and the fraction of reads mapping to organellar genomes. Red dashed lines indicate the filtering thresholds applied to each metric.

**Fig. S3.**
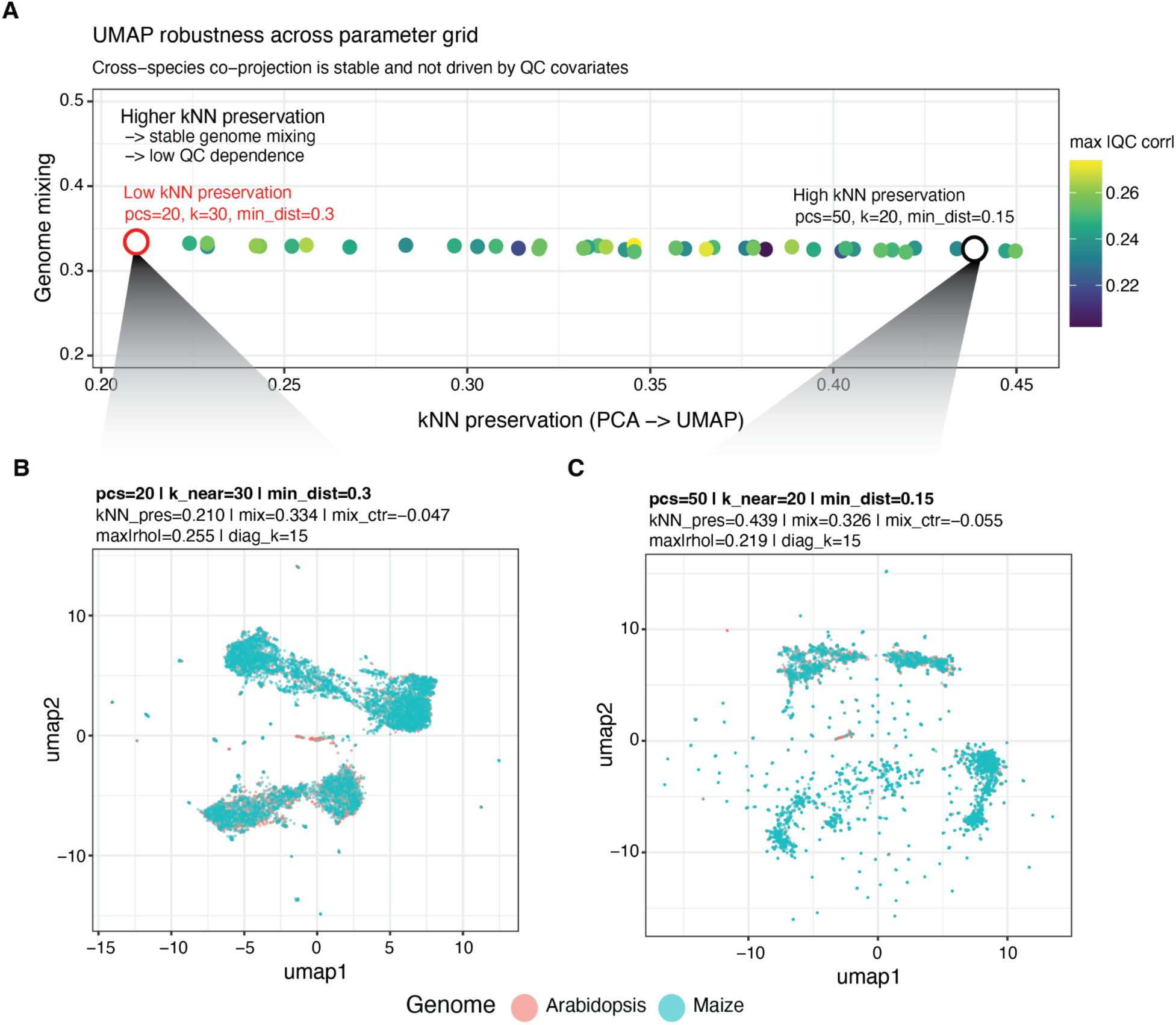
Cross species co-projection is robust to UMAP parameterization. **A,** UMAP performance across 45 combinations of component number, nearest neighbor number, and minimum distance using the concatenated reference object. Each point represents one embedding and is positioned by nearest neighbor preservation between PCA and UMAP space on the x axis and genome mixing on the y axis. Point color denotes the maximum absolute correlation between UMAP coordinates and the quality control covariates log10 nSites and pOrg. The two circled configurations are shown in **B** and **C**. **B,** UMAP generated using the parameter set applied throughout the study, with 20 components, k = 30, and min_dist = 0.30. **C,** UMAP generated using the parameter set maximizing the combined score for nearest neighbor preservation, centered genome mixing, and quality control independence, with 50 components, k = 20, and min_dist = 0.15. In both configurations, *Arabidopsis* and maize barcodes remain intermingled, indicating that the observed co projection is not driven by UMAP parameter choice.

**Fig. S4.**
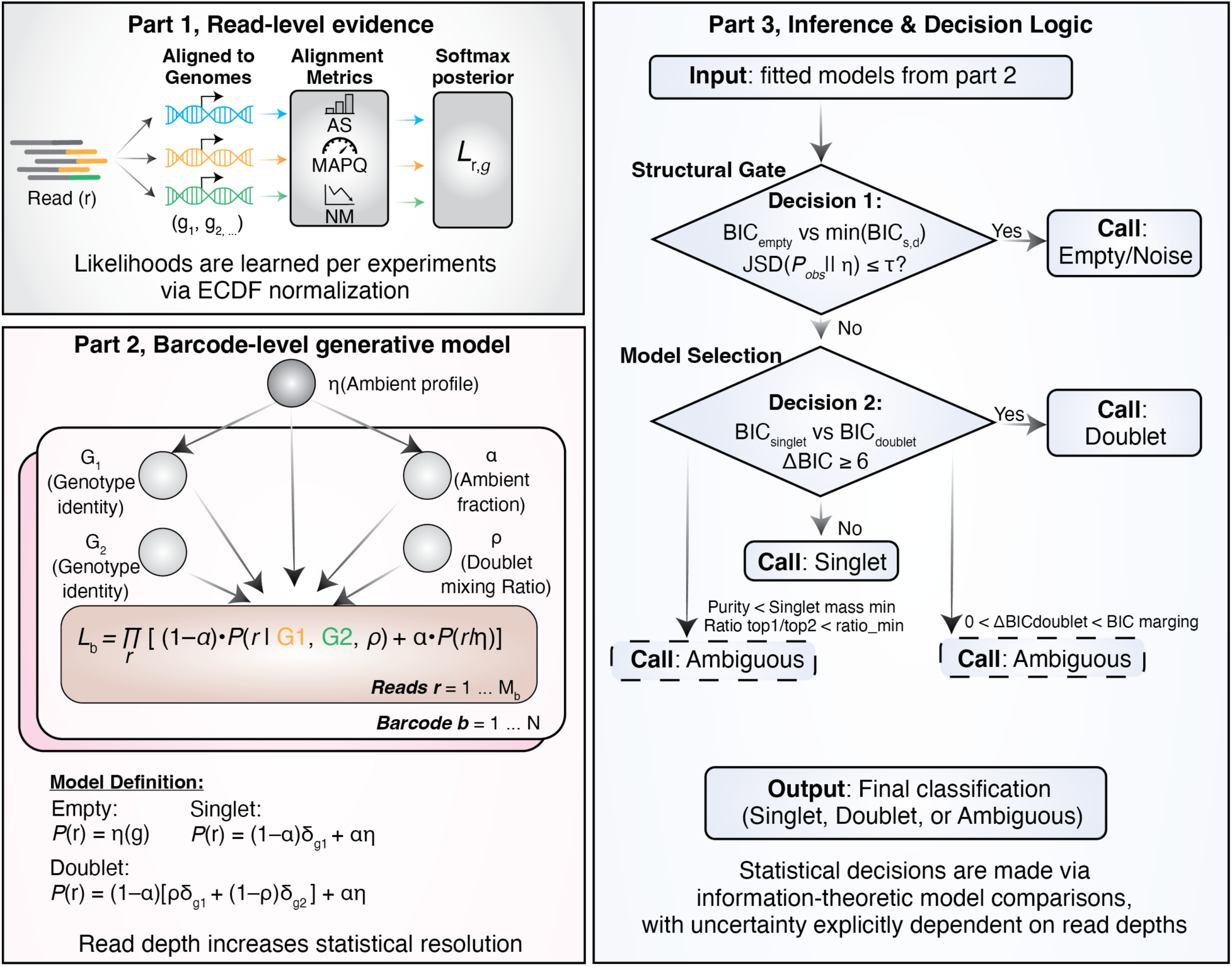
AmbientMapper model and decision calibration. Extended schematic of the AmbientMapper generative model and inference workflow. Read level likelihoods are derived from competitive alignment across candidate genomes and combined into barcode level likelihoods under empty, singlet, and doublet hypotheses, with statistical resolution increasing with the number of informative reads. In the first decision stage, barcodes consistent with the ambient profile are classified as empty when both the BIC criterion and ambient similarity criterion are satisfied. Barcodes passing this gate proceed to singlet versus doublet model selection using ΔBIC. Additional purity and strength criteria refine singlet and doublet assignments when the evidence is insufficient for a confident call. The final output assigns each barcode to singlet, doublet, ambiguous, or empty classes, with uncertainty explicitly dependent on read depth.

**Fig. S5.**
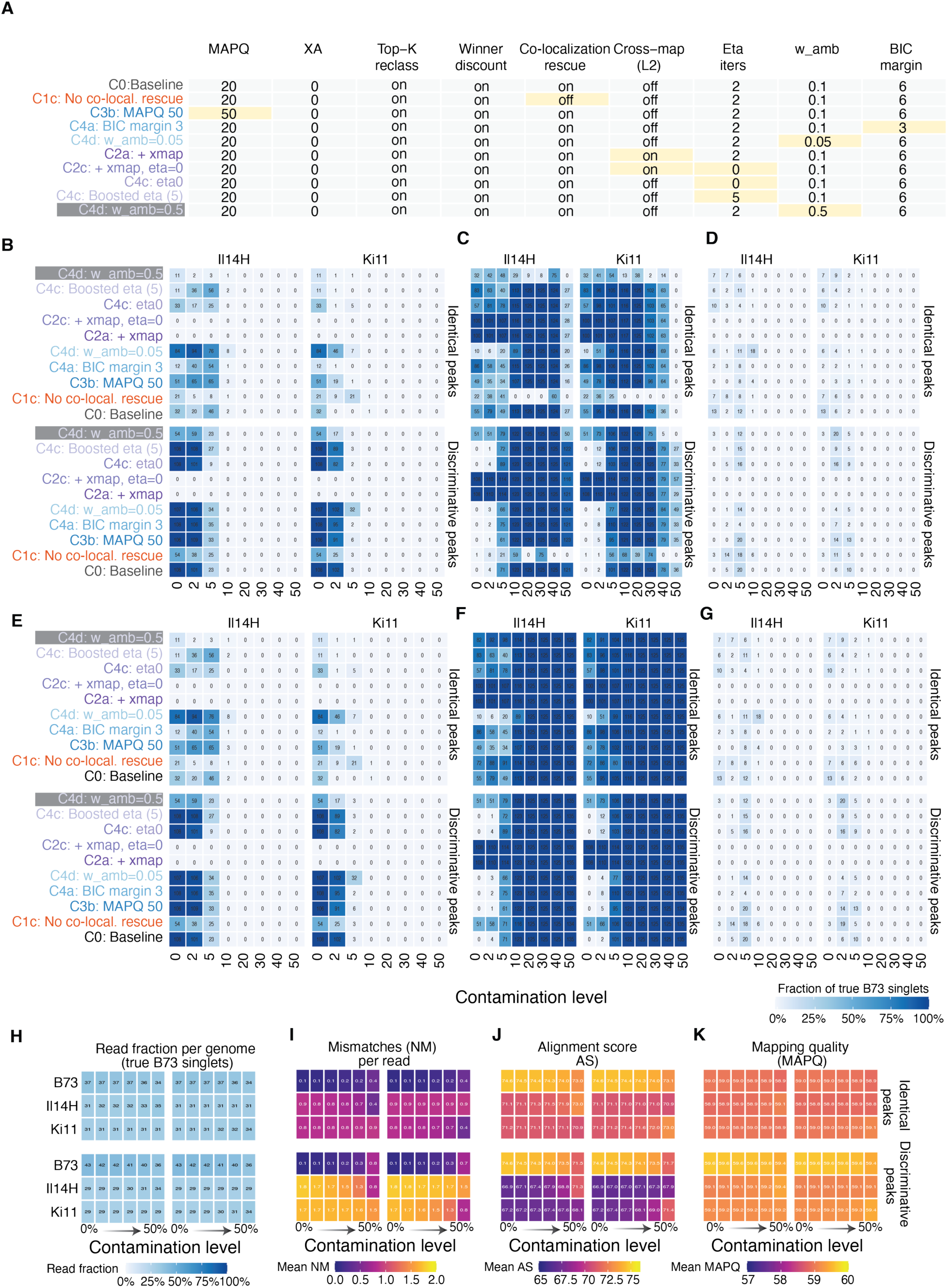
Genotyping parameter validation using a synthetic benchmark. All panels evaluate true B73 singlet templates with at least 500 reads across contamination levels α, separately for contaminant genome, Il14H or Ki11, and marker track. The identical peaks track contains ∼4000 accessible chromatin regions mappable one to one across B73, Il14H, and Ki11. The discriminative peaks track contains ∼1000 regions in which a 75 bp read is expected to overlap at least one variant distinguishing B73 from both Il14H and Ki11. **A,** Parameter configurations evaluated in the benchmark. Yellow tiles indicate settings that differ from the C0 baseline. **B to D,** Fraction of true B73 singlets whose top assigned genome is B73 and that are classified as singlet (**B**), doublet (**C**), or ambiguous (**D**) across configurations and contamination levels. **E to G,** Fraction of true singlets assigned to each call class regardless of top genome. **H to K,** Read level diagnostics for true singlet reads, showing the fraction assigned to each genome (**H**), mean edit distance NM (**I**), mean alignment score AS (**J**), and mean mapping quality MAPQ (**K**) across contamination levels. Printed values indicate template counts in **B to G** and panel means in **H to K**. Configurations C4c_eta0 and C4c_eta5 differ from C0 only in the number of ambient profile refinement iterations, which showed no measurable improvement in these runs, so differences among these configurations provide an estimate of run-to-run variability.

**Fig. S6.**
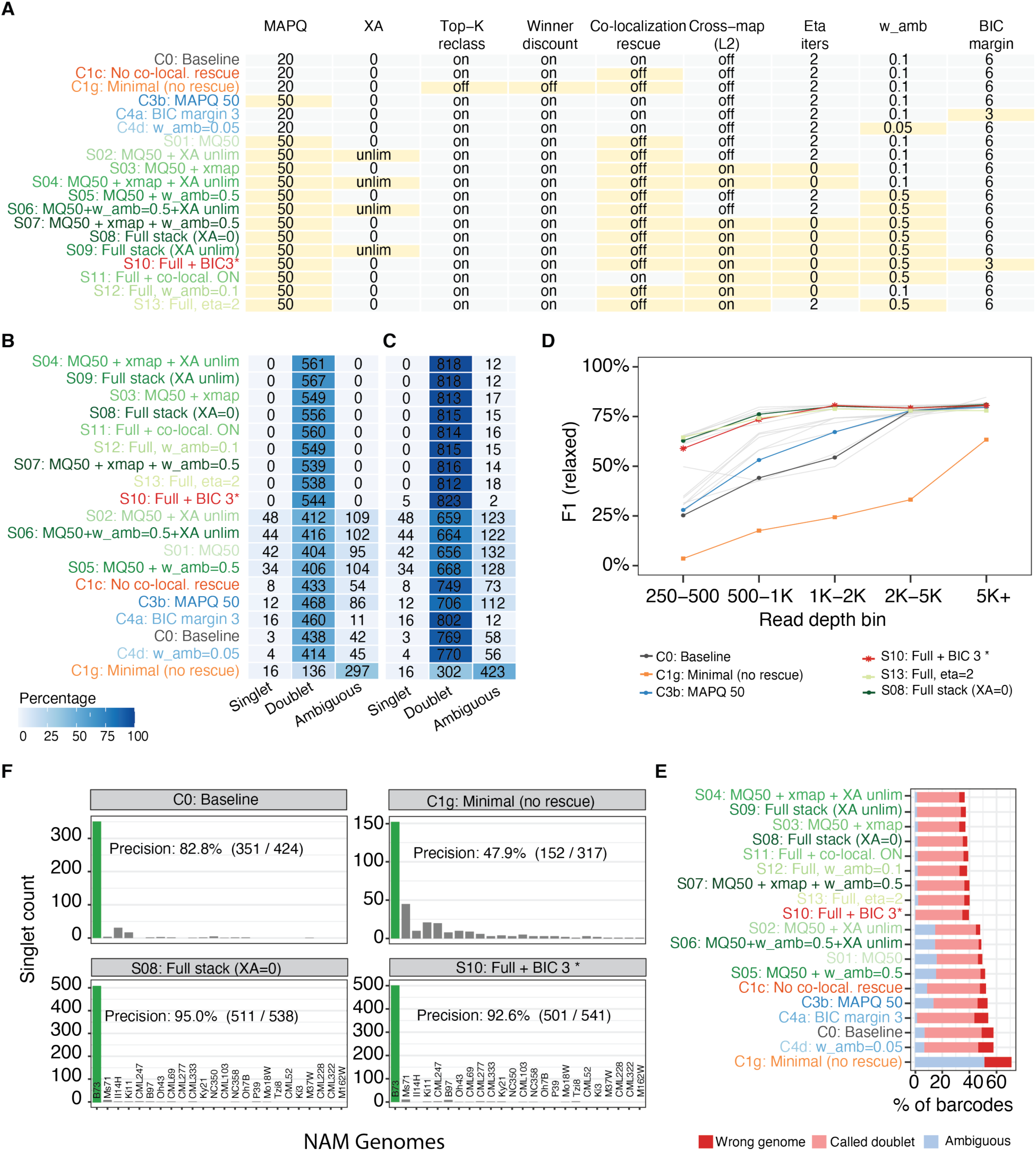
Genotyping parameter validation using a single genotype library. Nineteen AmbientMapper configurations were evaluated against a 26 genome NAM reference using a B73 only maize library, so any non B73 singlet assignment represents an error. All panels show the same library subsample, which reproduces the full library result. **A,** Parameter configurations evaluated in the benchmark. Yellow tiles indicate settings that differ from the C0 baseline. **B,** B73 dominance within each call class. Configurations are ordered by F1 score, with S10 ranked highest. Columns show singlet, doublet, and ambiguous classes, and labels indicate the number of barcodes in each class whose top assigned genome is B73. **C,** Barcode distribution across call classes regardless of top genome. Colors represent the fraction of barcodes with at least 500 nuclear reads assigned to each class. **D,** F1 score across read depth bins for all configurations. Six diagnostic configurations are highlighted and the remaining configurations are shown in gray. **E,** Failure modes across configurations, showing wrong genome, doublet, and ambiguous assignment rates, ordered by F1 score. **F,** Singlet genome assignments for four representative configurations, C0, C1g_naked, S08_full_xa0, and S10_full_bic3. B73 is the expected assignment, and calls to the remaining 25 NAM genomes represent errors.

**Fig. S7.**
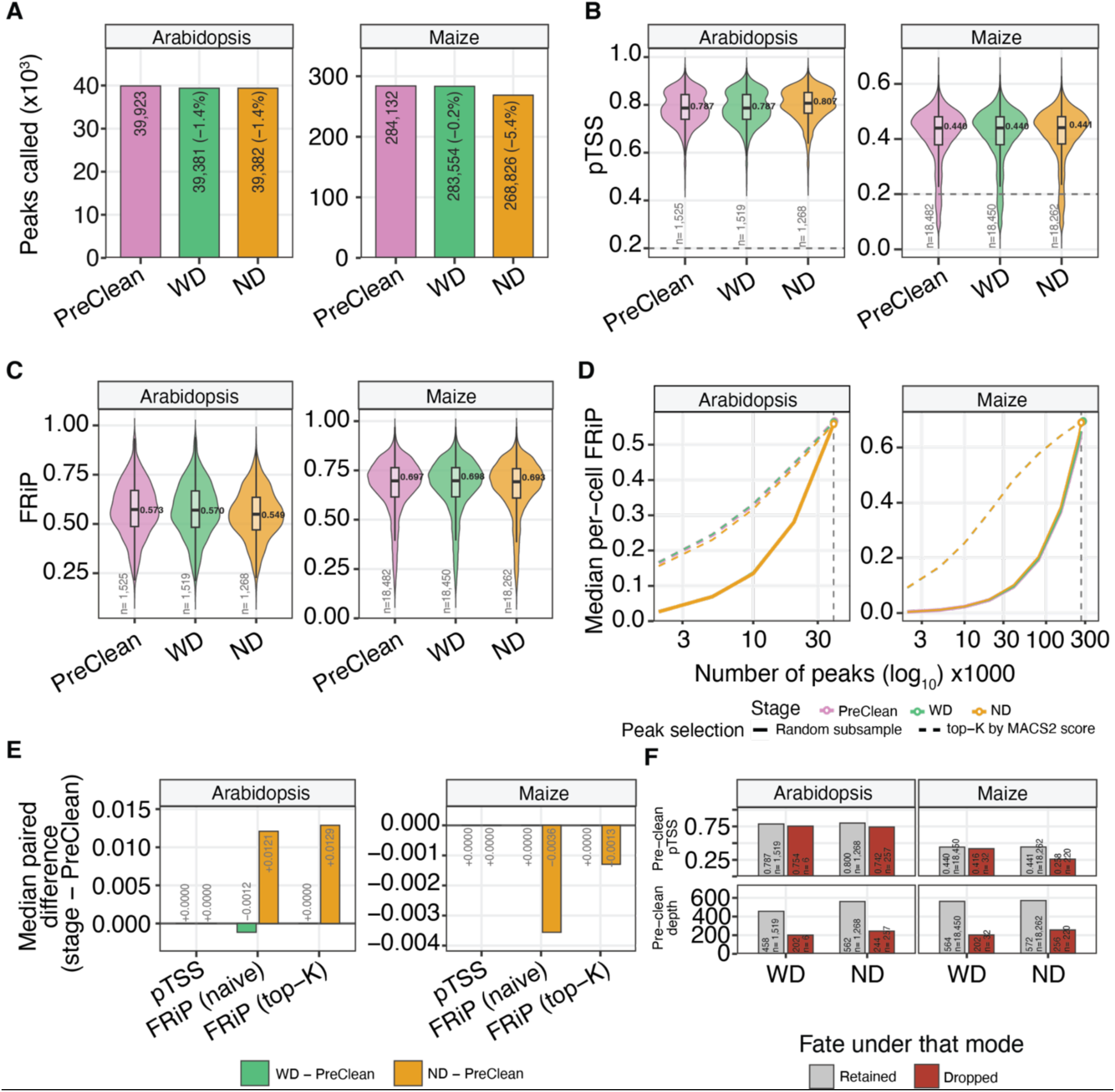
Cleaning preserves per cell data quality in the per genome analysis. All panels compare PreClean, WD, and ND stages for *Arabidopsis* and maize separately. **A,** MACS2 peak set size at each stage. The reduction in *Arabidopsis* peak number observed in the combined genome analysis is not reproduced in the per genome analysis. **B,** Per cell TSS enrichment across stages. Boxed values indicate medians, and the dashed line marks the 0.2 quality control threshold. Cells below this threshold are shown rather than filtered. **C,** Per cell FRiP calculated against the full peak set from each stage. **D,** Median per cell FRiP across matched peak counts using random peak subsampling and top K peak selection by score. Open circles indicate each stage using its complete peak set, and the vertical dashed line marks the common matched peak number. **E,** Median paired differences from PreClean for TSS enrichment, naive FRiP, and top K normalized FRiP. Y axis scales differ between genomes and should therefore be compared only within each panel. **F,** PreClean TSS enrichment and read depth for cells retained or removed by WD and ND cleaning.

**Fig. S8.**
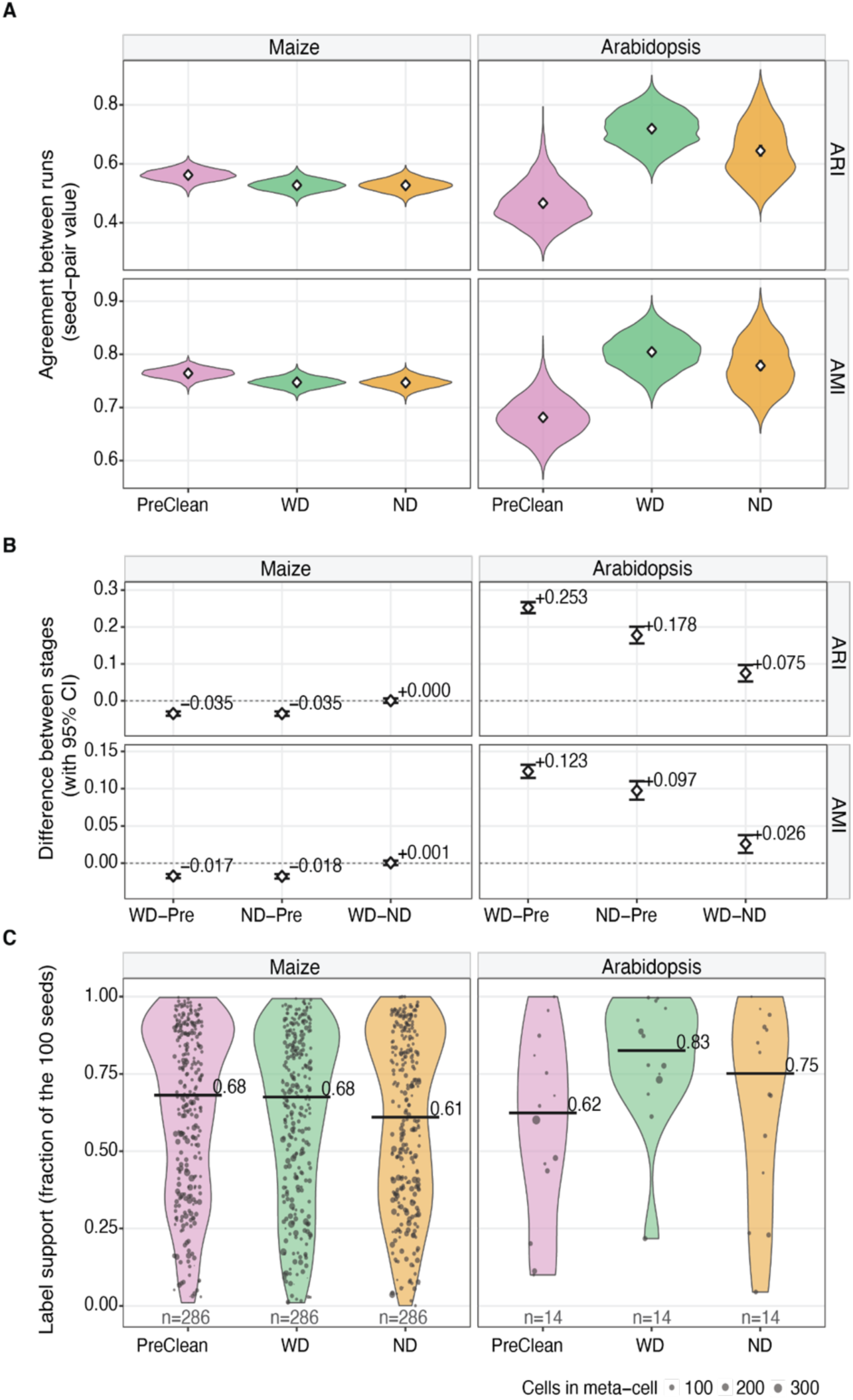
Meta cell partition agreement and annotation reproducibility across 100 SEACells runs. **A,** Pairwise agreement among meta cell partitions across 100 SEACells runs, quantified by adjusted Rand index, ARI, and adjusted mutual information, AMI. Each violin contains 4,950 pairwise comparisons. Values are descriptive because individual runs contribute to multiple pairwise comparisons. White diamonds indicate the mean, and bars show the 95% confidence interval obtained by bootstrap resampling of runs with 10,000 iterations. **B,** Differences in partition agreement between stages, with 95% percentile confidence intervals obtained from the same run level bootstrap procedure. **C,** Annotation reproducibility for each consensus meta cell, defined as the fraction of SEACells runs assigning the same cell type as the consensus annotation. Each point represents one consensus meta cell, with 286 maize and 14 *Arabidopsis* meta cells per stage. Point size indicates the number of cells per meta cell, and bars show the unweighted median for each stage. Values are descriptive because all meta cells derive from the same underlying partition and set of SEACells runs.

## Supplemental Tables

**Table S1 | Library and plate design of the scifi ATAC seq libraries analyzed in this study.** Each 96 well plate index identifies the well in which a nucleus was tagmented and, when a well contains a single sample, its sample of origin. This information was supplied to AmbientMapper in well design aware mode, WD. In the maize and *Arabidopsis* library, each species occupied a separate set of wells. The B73 and Mo17 library was pooled before tagmentation and therefore lacked genotype specific well information. In the multi genotype library, plate design was withheld from AmbientMapper and used only as independent ground truth. The maize root library was generated using 10x scATAC and contained no combinatorial plate design.

**Table S2 | Datasets and reference panels used in this study.** For each dataset, the table reports the candidate reference genomes used by AmbientMapper and whether plate design information was supplied during inference. Only the maize and *Arabidopsis* library was analyzed in both WD and no design, ND, modes. The multi genotype library was analyzed without design information, with its plate layout retained as independent ground truth. The synthetic benchmark provides exact genome of origin for every simulated read.

**Table S3 | Cross plate barcode misassignment by plate origin and read depth.** Analysis includes WD barcodes from the maize and *Arabidopsis* library classified as *singlet* or *dirty singlet* with at least 100 reads before cleaning. Wrong top genome indicates barcodes whose highest supported genome differs from the genome expected from the plate design. Pure wrong additionally requires less than 1% of reads to support the expected genome, consistent with barcode misassignment. The maize plate direction provides the least confounded estimate because *Arabidopsis* ambient chromatin is rare in maize indexed barcodes. The decline in the *Arabidopsis* plate rate with increasing depth is more consistent with ambient filling than with barcode level misassignment.

**Table S4 | Support for grouping weak doublet barcodes with singlets in the allele purity analysis.** Weak doublets favor the two-genome model by BIC but fail an additional strength criterion, indicating limited evidence for a true doublet. At genotype informative sites, these barcodes predominantly behave as singlets. In the multi genotype library, the dominant genome accounts for a median of 93% of reads assigned to the inferred genome pair, compared with 50% expected for an equal doublet. Their contribution is negligible in the B73 and Mo17 libraries but substantial in the multi genotype singlet group, and this difference should be considered when interpreting Fig. 4L.

